# Modulation of Neuronal Excitability and Plasticity by BHLHE41 Conveys Lithium Non-Responsiveness

**DOI:** 10.1101/2024.07.25.605130

**Authors:** Marius Stephan, Sergi Papiol, Mingyue Zhang, Jie Song, Samuel M. Frommeyer, Helen Haupt, Niels Jensen, Nirmal Kannaiyan, Rajinder Gupta, Philipp Schuler, Pia Picklmann, Michael McCarthy, Eva Schulte, Mikael Landen, Peter Falkai, Volker Scheuss, Thomas Schulze, Weiqi Zhang, Moritz J. Rossner

## Abstract

Many bipolar disorder (BD) patients are non-responsive to lithium. The mechanisms underlying lithium (non-)responsiveness are largely unknown. By using gene-set enrichment analysis methods, we found that core clock gene-sets are significantly associated with lithium response. Among the top hits was *BHLHE41*, a modulator of the molecular clock and homeostatic sleep. Since BHLHE41 and its paralog BHLHE40 are functionally redundant, we assessed chronic lithium response in double-knockout mutant mice (DKO). We demonstrated that DKOs are non-responsive to lithium’s effect in various behavioral tasks. Cellular assays and patch clamp recordings revealed lowered excitability and reduced lithium-response in prefrontal cortical layer 2/3 DKO neurons and on hippocampal long-term potentiation. Single-cell RNA sequencing identified that lithium deregulated mitochondrial respiration, cation channel and postsynapse associated gene-sets specifically in upper layer excitatory neurons. Our findings show that lithium acts in a highly cell-specific way on neuronal metabolism and excitability and modulates synaptic plasticity depending on BHLHE40/41.

## Introduction

Bipolar Disorder (BD) is one of the most prevalent disorders with a high global disease burden and BD patients have a 20 to 30-fold increased suicide rate ^1^. Symptomatically, BD is primarily a mood disorder characterized by recurrent episodes of mania and depression ^2^. Lithium, in the form of its salts, is a first line medication for prophylaxis of episodes and reduction of suicidality in BD ^3^. Lithium is unique among neuropharmacological compounds as a simple alkaline metal ion and has a long and rich history in the treatment of neuropsychiatric conditions. Lithium interacts with numerous proteins, making the distinction between therapy-relevant and side or even toxic effects difficult ^4^. Elucidating lithium’s mode of action is of particular interest because of a major unmet clinical need: only about a third of BD patients are considered excellent responders, while remaining patients show only partial to no response to lithium treatment ^5^.

A deeper understanding of the therapeutic mechanism of lithium and the factors that contribute to non-responsiveness might lead to new treatments with the beneficial properties of lithium, but higher efficacy in resistant patients and a decreased risk of side effects. Reliable mouse models for lithium non-responsiveness, however, are lacking. In the past, many researchers have linked new findings on BD pathogenesis to mechanisms of lithium response and vice versa, assuming an overlapping genetic and neurophysiological basis ^6,7^. However, although there is undeniably a mechanistic link between pathophysiology and recovery, the distinction between trait and state is key in the treatment of psychiatric disorders as the neurodevelopmental pathophysiology might not be reversible. Therefore, effective therapies might have to rely on the modulation of the neuronal function and residual plasticity present in the adult brain^10^.

Along these lines, lithium’s effect on neuronal excitability and its neuroprotective properties were extensively studied in the past. Compelling evidence for excitability-dampening as one of lithium’s main therapeutic effects was demonstrated with patient-derived neuronal cultures ^11^. Neurons generated from lithium responders showed reduced neuronal hyperexcitability upon lithium treatment which was not observed in cultures derived from lithium non-responder patients ^11^. More recently, it has been shown that lithium increases mitochondrial respiration in patient-derived neural precursor cells from lithium responders and partially rescued excitability in BD patient derived cortical spheroids ^12,13^. These findings suggest that lithium (non-)responsiveness is, at least partially, cell autonomous. In addition, it clearly demonstrates a genetic basis of lithium (non-)responsiveness. This is supported by studies showing that lithium response is a familial trait ^14^, so we can assume that there is not only a major contribution of genetics to BD risk but also to (non)responsiveness to lithium treatment and that the genetic architecture of lithium (non-)responsiveness likely represents a distinct trait that may partially overlap with BD risk.

The molecular clock represents a possible mechanism of convergence between BD and lithium response. BD patients were reported to exhibit abnormalities in circadian rhythms, with alterations in the sleep/wake cycle during manic and depressive phases and sleep problems during euthymia ^15,16^. Moreover, mouse models for modulators of the circadian clock genes exhibit mood related ^17^ and mixed-state endophenotypes of BD ^18–20^.

The circadian clock is a central regulator of neuronal redox metabolism and excitability in the adult mammalian brain in the suprachiasmatic nucleus (SCN) ^21,22^ and hippocampus (HIP) ^23^. This is supported by evidence that cortical excitability underlies circadian regulation in humans ^24^ and that circadian change of membrane potential regulates sleep states ^25^. Taken together, circadian modulation of neuronal excitability is a promising mechanism to be involved in BD symptoms and recovery mechanisms.

Despite some suggestive results, the implication of clock genes as risk factors for BD has not yet been firmly established in humans. The strongest association of a core clock gene with BD was obtained for *ARNTL* (or *BMAL1*) ^26^, which is the only non-redundant clock gene ^27^. One potential explanation for these inconclusive results is that genetic association studies primarily aim to estimate the individual effect of each genetic variant rather than analyzing genetic variants in functionally connected sets of genes. Gene-set analysis holds promise to overcome this limitation by integrating the polygenic architecture of these traits with biological information ^28^. The molecular clock represents a prototypical set of functionally connected genes, whereby components of the ‘core clock’ are implicated in the circadian translational-transcriptional feedback loop and ‘clock modulator or output’ genes are thought to properly align metabolic and homeostatic processes in an organ and cell-type specific fashion to the circadian cycle ^27^.

In this study, we have defined the following gene-sets related to the circadian system, based on previous empirical evidence: 1) core clock genes 2) clock modulator genes, and 3) genes with a circadian pattern of expression. We use GWAS summary statistics of major psychiatric disorders, chronotype, and lithium response as the basis for gene-set association analyses. We detected a significant enrichment of core clock gene-sets in lithium response in BD using two different analysis methods ^29,30^ and replicated our findings in an independent GWAS on lithium response. We identified BHLHE41 (also known as SHARP1 and DEC2) as promising candidate for further investigation and show that *Bhlhe40/41*^−/−^ double-knockout (DKO) mice are non-responsive to lithium. We observed reduced excitability in DKO cortical neurons in culture as well as glutamatergic neurons in layer 2/3 of the anterior cingulate cortex (ACC) and the CA1 region of the ventral hippocampus (vHIP). A lithium dependent, cation channel and postsynapse-dominated gene-set was found to be specifically deregulated in layer 2/3 pyramidal neurons of the prefrontal cortex. Furthermore, DKO neurons were non-responsive in chronic lithium treatment-induced excitability dampening. *Arntl* heterozygous mice were lithium responsive, and we show that *ARNTL* is associated with BD but not lithium-response. Taken together, our analyses thus show that components of the circadian clock are associated with different aspects of BD including control of neuronal excitability and plasticity, and treatment response.

## Results

### The core clock gene-set including *BLHE41* is strongly associated with the lithium response

MAGMA analyses for 13 preselected traits revealed the significant association of the core clock gene-set (competitive P_corrected =_ 0.0046) with lithium response on a dichotomous definition (responder/non-responder) of the Alda scale ^14^ (Consortium on Lithium Genetics [ConLiGen] GWAS) (Figure 1A). No other statistically significant gene-set enrichments were observed in any of the other psychiatric disease traits analyzed, including bipolar disorder (Figure 1A). Two core clock gene-sets and one circadian prefrontal cortex associated gene-set reached significance with the chronotype GWAS data, validating the approach (Figure 1A). INRICH analyses performed with 3 core clock gene-sets revealed similar results, showing a significant association with chronotype GWAS at all p-value thresholds and with lithium response at lowered p-value thresholds (Figure S1A). MAGMA and INRICH analyses of the independent Swedish cohort revealed significant associations of core clock gene-sets on the subjective scale used for the assessment of lithium response in this study (Figure S1B, C).

**Figure 1.**
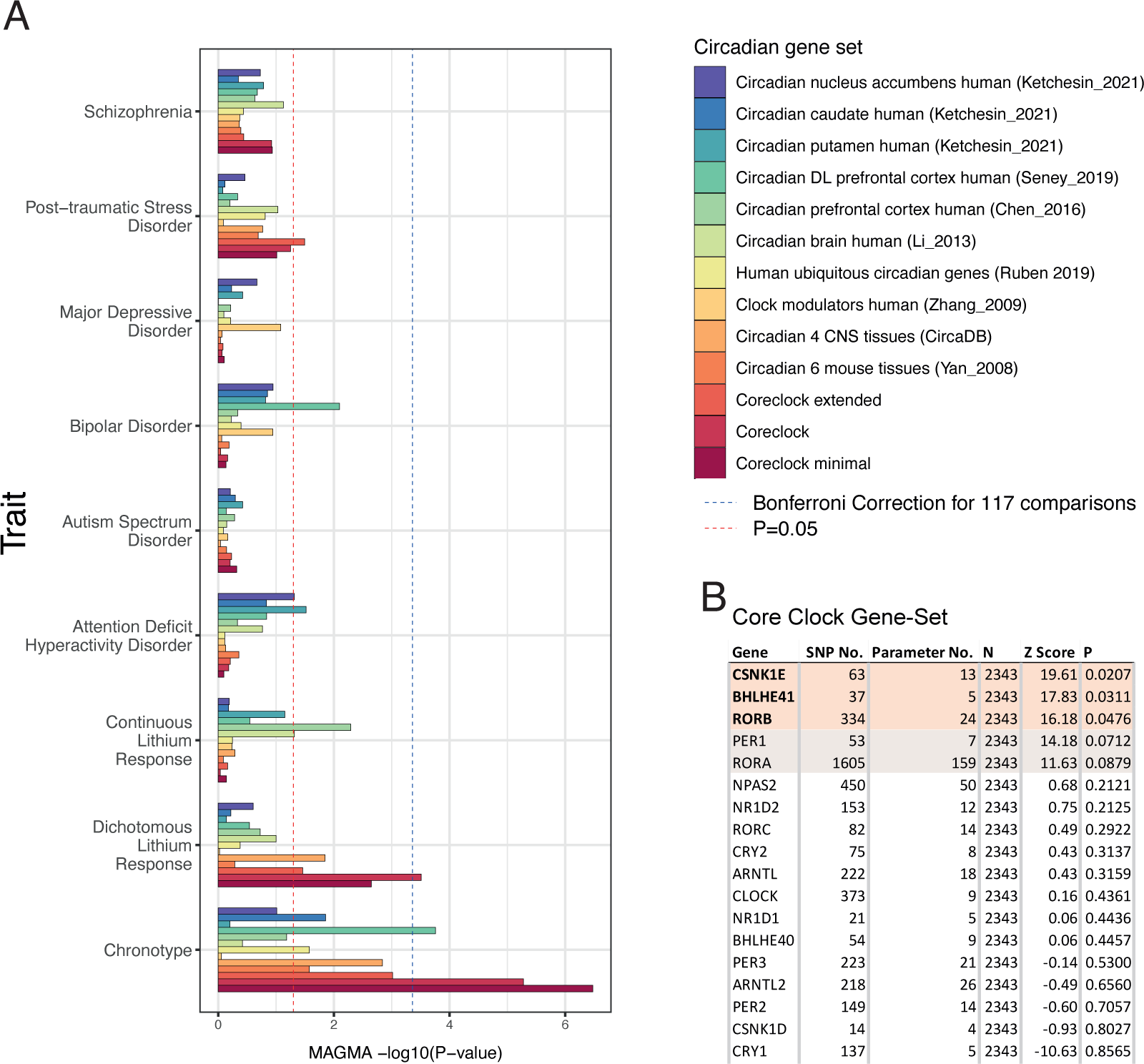
BHLHE41 is a strong contributor to the association between lithium response and the core clock geneset. (A) MAGMA analyses based on circadian gene-sets in psychiatric traits. Competitive test P-values in the X-axis are –log10 converted. Dashed blue line indicates the Bonferroni significance threshold (P<0.05/8 gene-sets x 9 traits). Dashed red line indicates the nominal (uncorrected) significance threshold P=0.05. (B) In gene-based tests three genes reach nominal significance (unadjusted p-value). (C-E) Local Manhattan plots of these genes (+/-15 KB) for dichotomous lithium response show the BHLHE41 locus (C) has the strongest association (5 SNPs with r² > 0.8) compared to CSNK1E (D) and RORB (E). ADHD: Attention-Deficit/Hyperactivity Disorder; PTSD: Post-Traumatic Stress Disorder.

Using MAGMA gene-based tests on the individual member genes of lithium response-associated core clock gene-sets, the genes *RORB*, *BHLHE41*, and *CSNK1E* reached nominal significance (uncorrected P < 0.05) and the genes *PER1* (P = 0.0712) and *RORA* (P = 0.0879) were close to significance. These genes, except for *PER1*, are not components of the core transcriptional-translational loop (TTL) (Figure 1B), suggesting that output genes of the molecular clock are the major contributors of this association. The genome-wide gene-based association study (GWGAS) showed that *BHLHE41* has the strongest association of the three top genes (Figure S2). In contrast, no such association for *BHLHE41* and *CSNK1E* was found for BD life-time risk, in contrast to the strong signal obtained for *RORB* and *ARNTL* (Figure S2). Of note, BD risk was also not enriched in the MAGMA analyses (Figure 1A), suggesting that *BHLHE41* (which was also identified with INRICH at the gene level (Table S1) and *CSNK1E* are primarily associated with treatment response and neither with pathogenesis of BD nor any other psychiatric disease risk studied.

Taken together, our genetic analysis with independent methods and cohorts indicates a central role of BHLHE41 in the regulation of disease course and severity in response to lithium treatment.

### Bhlhe40/41^−/−^ mutant mice are resistant to lithium’s effect on cognition

After identifying *BHLHE41* as a major contributor to the genetic association of circadian clock genes and lithium response in patients, we used *Bhlhe40/41^−/−^* double-knockout (DKO) mice as a genetic model of altered lithium responsiveness. We focused on DKO mice because of the functional redundancy of *Bhlhe40* and *Bhlhe41* in circadian entrainment, light adaptation phenotypes ^31^ and cognitive performance ^32^.

Therefore, we conducted a behavioral study using our standardized platform for systematic cognitive and behavioral profiling (PsyCoP) ^33,34^ to assess the neuro-cognitive profile of chronically lithium-treated DKO and WT mice (Figure 2A, Table S2). Drinking behavior during the IntelliCage period was not significantly different between genotypes as well as treatment groups (Figure S3A). We detected a significant statistical interaction of genotype and treatment (GxT) in the overall PsyCoP profile (MANOVA G: P = 2.98 × 10^−6^, T: P = 7.79 × 10^−7^, GxT: P = 0.0278, Table S3; normality controls shown in Figure S3B – C). The circadian 24h activity profile was mainly altered at peak activity phases in D (ZT13-20), here we detected a significantly reduced activity in DKOs (G: p<0.05) and strong reduction upon lithium treatment in both genotypes (T: p<0.001), and a trend for GxT interaction (p and in several aspects of the cognitive systems domain (Figure 2C – E). Contextual fear memory was improved in lithium treated (LiCl) WT mice compared to vehicle control (Ctrl) but not in DKO mice, in recent (24 hours; Figure 2B; Context: GxT: P_adj_ = 0.0464; WT-T: P = 0.0165; DKO-T: P = 0.942) and remote tests (21 days; Figure 2C; Remote: GxT: P_adj_ = 0.0464; WT-T: P = 0.00705; DKO-T: P = 0.767). Similarly, in the serial reversal test of spatial learning, WT lithium treated mice needed less trials to reach the learning criterion, and therefore learned faster than WT Ctrl mice, especially in the first two reversal phases, resulting in significantly smaller area under the learning curve (Figure 2D, E). In contrast, serial reversal learning performance did not improve in DKO mice, resulting in a strong interaction of genotype and treatment, indicating non-responsiveness of DKO mice to the chronic lithium treatment effect (SerialLearn: GxT: P_adj_ = 0.00248; WT-T: P = 0.00675; DKO-T: P = 0.0857).

**Figure 2.**
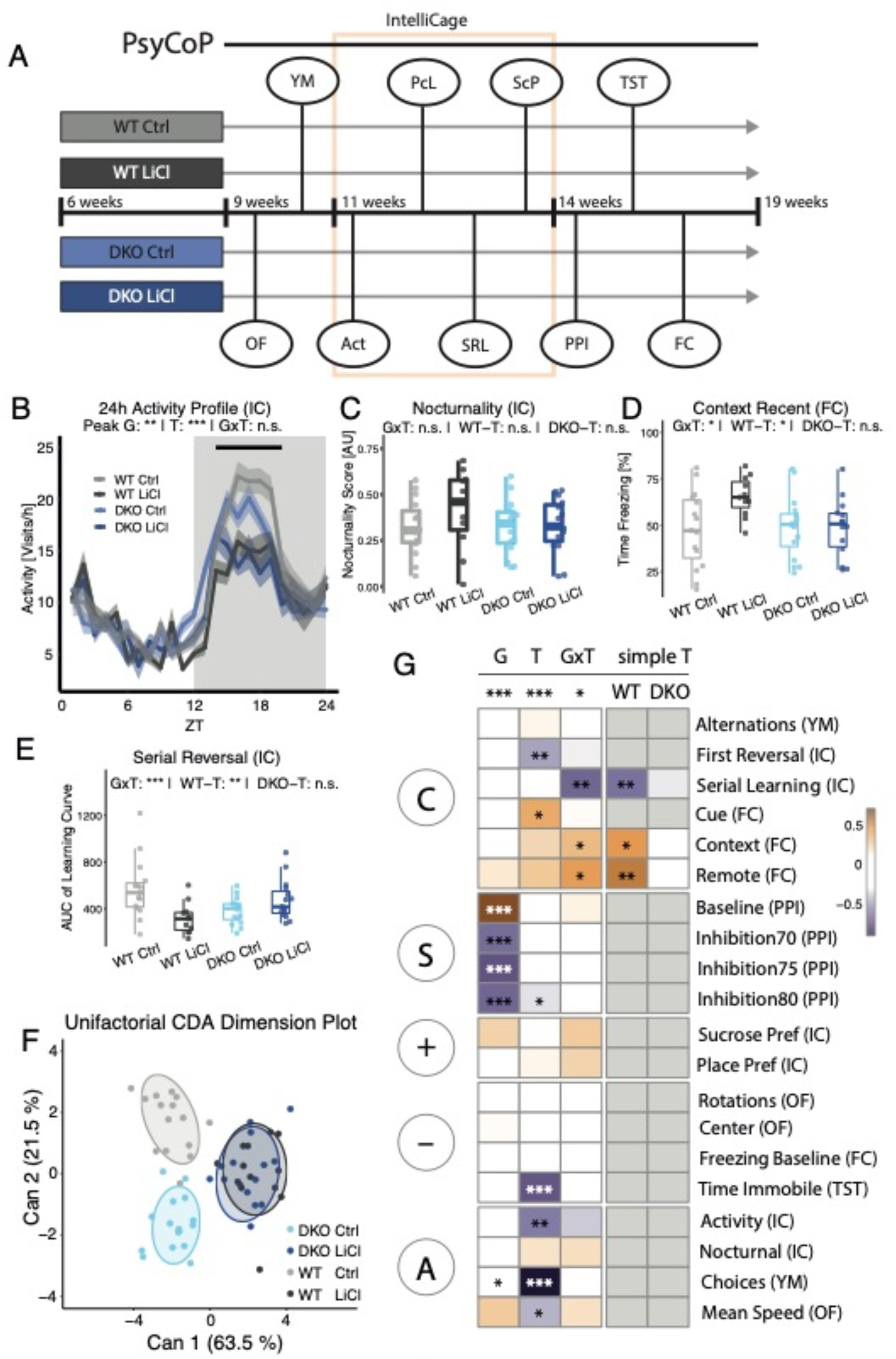
PsyCoP profile reveals resistance of BHLHE40^-/-^/41^-/-^ mice to lithium’s beneficial effect on memory and cognition. (A) Schematic of the experiment design. Four groups, two from each genotype and sex, were tested after three weeks of pre-treatment via with either Vehicle (Ctrl) or lithium chloride solution (LiCl). Treatment was continued throughout the experiments. (B) Circadian 24h activity plotted as group means of 1-hour bins and ribbons show the standard error of the mean. The light and dark bars visualize the light and dark phase, respectively. Black line indicates time points used to assess peak activity differences revealing significant genotype (G* = p < 0.05) and lithium treatment (T *** p < 0.001) dependent effects, as indicated. (C-E) Boxplots showing lithium-affected phenotypes in DKO animals. Data are shown as box plots with whiskers extending to no more than 1.5-fold IQR. (C) Nocturnality (relative activity differences between L and D phases) revealed a non-significant tendency of Lithium non-response in DKOs. Lithium treatment resistance was found to be significant in DKOs in (D) recent contextual fear memory and (D) Serial Reversal Learning performance, as indicated. (F) Dimension reduction showing the canonical scores of the first two canonical components Can1 and Can2 (explaining 63.5% and 21.5% of the overall canonical correlation, respectively) depicts the aggregated Li-treatment dependent difference in WT but not DKO animals. Datapoints are overlayed with a 75% data ellipse. (G) A heatmap of the weights of single variables in the structure of the CDA for each term of the profile’s linear model. The variables are grouped in RDoC Top Level Domains as assigned *a priori*. Multivariate ANOVA results are shown on top. Univariate contrasts reaching significance are indicated with asterisks in the respective panel.* p < 0.05; ** p < 0.01; *** p < 0.001; n.s. not significant; p-values are FDR-adjusted and refer to Wilk’s lambda testing two-way ANOVA; simple effects were tested in a similar but univariate and unifactorial ANOVA procedure; WT: wildtype mice; DKO: BHLHE40^-/-^/41^-/-^ mice; Ctrl: vehicle control; LiCl: lithium-treated; OF: Open Field Test; YM: Y-Maze Test; IC: IntelliCage; Act: circadian activity; PcL: Place Learning; SRL: Serial Reversal Learning; ScP: Sucrose Preference Test; PPI: Prepulse Inhibition Test; TST: Tail Suspension Test; FC: Fear Conditioning Test; G: genotype main effect; T: treatment main effect; GxT: interaction effect; simple T: simple effects; DKO-T: DKO simple treatment effect; WT-T: WT simple treatment effect; C: Cognitive Systems Domain; S: Sensorimotor Systems Domain; “+”: Positive Valence Systems Domain; “-“: Negative Valence Systems; A: Arousal and Regulatory Systems Domain

After CDA dimension reduction (Figure 2F), we observed that ellipsoids of vehicle-treated (Ctrl) DKO mice are clearly different from their wildtype counterpart, whereas lithium treated groups are completely overlapping indicating lowered response to treatment. When looking at the variable weights (Table S2), it becomes apparent that the shift is mainly driven by the modulation of cognitive aspects in WT mice. A heatmap of variable contributions to the genotype (G), treatment (T), and interaction effects (GxT) shows the domain-specific character of the non-response to lithium in DKO mice (Figure 2G). In the cognitive domain, we find significant interaction for contextual fear memory and serial reversal learning, but not for cue-dependent fear memory (Table S3 and Figure S4; Cue: GxT: P_adj_ = 0.284). This indicates that the observed non-responsiveness may depend primarily on the medial prefrontal cortex and ventral hippocampus but not on the basolateral amygdala ^35^.

Sensorimotor gating, on the other hand, was almost exclusively influenced by genotype, with only a minor lithium effect (9.5 % mean decrease) at a prepulse level of 80 dBA (Table S3, Figure S4D – G; Inhibition80: T: P_adj_ = 0.371). We did not observe lithium treatment effects in positive valence (Table S3 and Figure S5) but a strong treatment effect for the time spent immobile in the tail suspension test (TimeImmobile: T: P_adj_ = 4.44 × 10^−6^) in the negative valence domain. TimeImmobile is possibly influenced by the general activity level, since arousal and regulatory domains showed a strong treatment dependent reduction in both novelty-induced locomotor activity in the open field (MeanSpeed: G: P_adj_ =0.207, T: P_adj_ = 0.0161, GxT: P_adj_ = 0.298) and Y-maze (Choices: G: P_adj_ = 0.0317, T: P_adj_ = 5.82×10^-13^, GxT: P_adj_ = 0.437) as well as general locomotor activity in the IntelliCage (Activity: G: P_adj_ = 0.0734, T: P_adj_ = 0.00152, GxT: P_adj_ = 0.145), independent of genotype (Figure S6). The only clear sex-dependent treatment effect was found for general activity, where males showed a stronger reduction in activity level than females (Table S3 and Figure S7; TxS: P_adj_ = 0.0373).

Based on these results and the previous observation that *ARNTL* at the gene-level was associated with BD but not the lithium response (Figure 1B, Figure S2), we analyzed the lithium response in *Arntl* heterozygous null mutants (Het). *Arntl* Het mice were chosen since these animals display psychiatric endophenotypes ^36^ and homozygous mutant animals display enhanced postnatal mortality and are lacking any circadian rhythm ^37^. In contrast to DKOs, *Arntl* Hets and control mice showed a non-significant dampening of the baseline activity by lithium and a significant lithium response in the contextual fear memory task (Figure S8). As with DKOs, cue fear memory remained unchanged, arguing for a hippocampal but not amygdala dependent effect (see above).

In summary, *Bhlhe40/41* DKO mice were non-responsive and *Arntl* Hets responded normal to lithium treatment in the cognitive domain, most evident in tasks associated with hippocampal and cortical functions. These translational observations support the assumption that BHLHE40/41 are modulators of the lithium response but not ARNTL.

### Cortical neurons of DKO mice are less responsive to lithium treatment

Lithium has been shown to reduce excitability in lithium-responsive patient-derived neurons, but not in neurons derived from lithium non-responsive patients ^11^. Moreover, mRNA expression of *Bhlhe41* and *Bhlhe40* is induced by neuronal activity ^38^, and BHLHE40 has previously been associated with control of neuronal excitability ^39,40^. To test excitability in primary neuronal cultures, we first applied a transcriptional reporter system for neuronal network activity the ‘enhanced synaptic activity response element’ (ESARE), that measures synapse-to-nucleus signaling ^41^ (Figure 3A).

**Figure 3:**
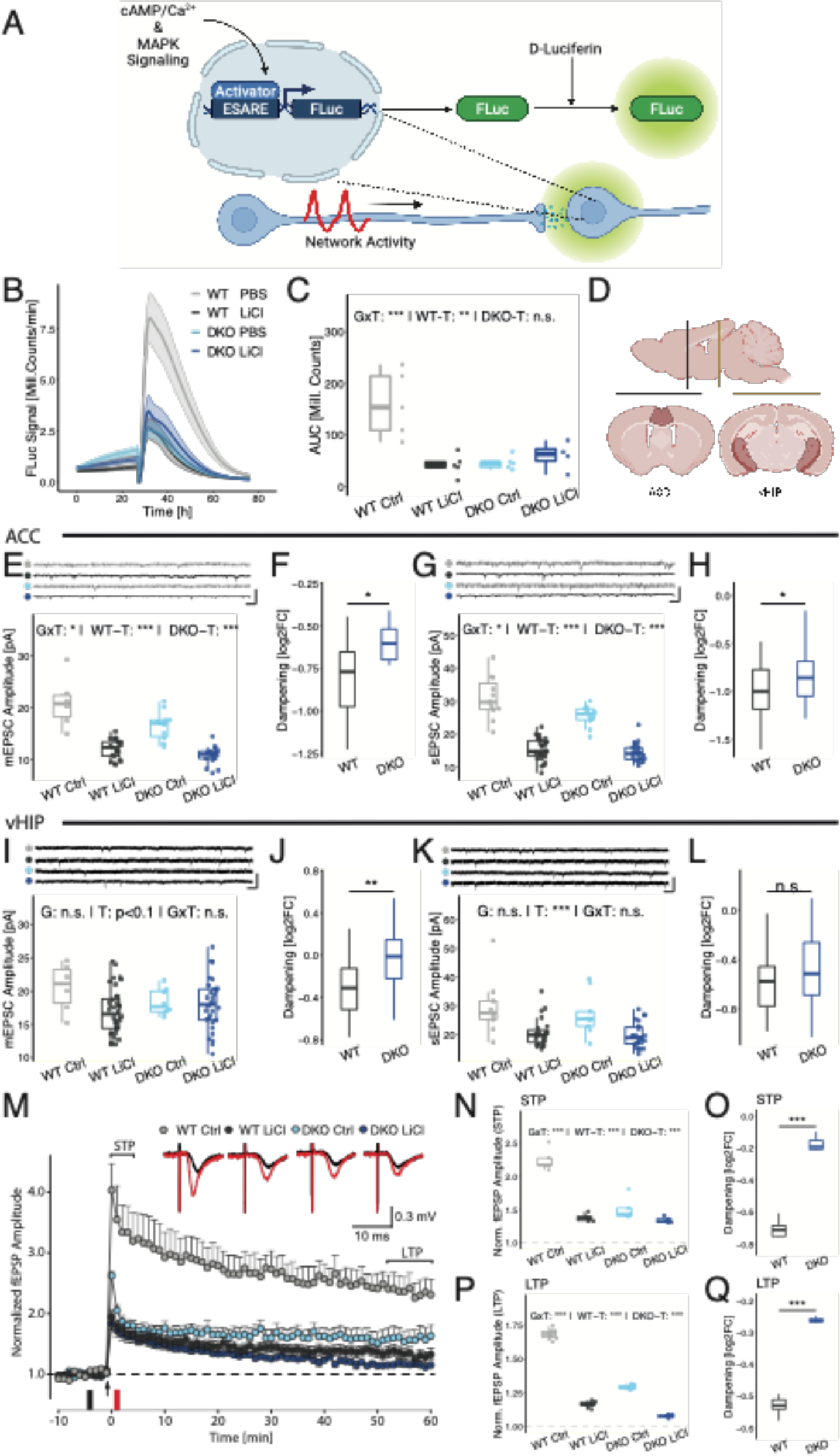
Primary cortical neuron cultures and hippocampal networks in BHLHE40^-/-^/41^-/-^ mice are resistant to lithium’s stimulation dampening effect. (A) To measure neuronal network activity in primary cortical neuron cultures, a previously published^68^ Firefly luciferase reporter assay was used. This assay is based on a sensor for synapse-to-nucleus signaling named “enhanced synaptic activity response element (ESARE). (B) Ribbon plots showing the mean ESARE activity in response to bicuculline (BIC) stimulation after 24 hours. The ribbons indicate the standard error of the mean. (C) Quantification of the stimulation peak. Data are shown as box plots with whiskers extending to no more than 1.5-fold IQR. Asterisks indicate statistical significance of the interaction effect and simple effects for both genotypes tested in a two-way ANOVA. (D) The regions of interest for electrophysiology in acute slices were the anterior cingulate cortex (ACC) portion of the medial prefrontal cortex (mPFC) and the ventral Hippocampus (vHIP). (E) Shows the exemplary placement of the patch pipette in layer 2/3. Excitatory Postsynaptic Currents (EPSCs) were quantified in mPFC (F-I) and vHIP (J-M). (F, H) miniature EPSC (mEPSC) amplitude (F) and spontaneous EPSC (sEPSC) amplitude (H) in L2/3 mPFC excitatory neurons were significantly dampened in both WT and DKO, but the significant interaction of genotype and treatment (GxT) in both variables indicates a difference in lithium-induced dampening. (G, I) A significantly smaller log2-transformed foldchange (log2FC) reduction in LiCl-treated compared to Ctrl-treated neurons in both mEPSC (G) and sEPSC (I) amplitude supports this finding. (J) Average mEPSC amplitude was not significantly reduced in CA1 principal neurons. (K) However, the nominal reduction in amplitude is significantly stronger in WT compared to DKO. (L) The average sEPSC amplitude is reduced in both WT and DKO CA1 neurons in response to lithium treatment. (K) Normalized to the mean amplitude in Ctrl treated mice, we find a significantly stronger dampening effect in WT neurons compared to DKO. (N) Short-term (STP) and long-term potentiation (LTP) was assessed by high frequency stimulation (HFS) of the Schaffer collaterals and measuring fEPSP in the CA1 region of the hippocampus. Time point of stimulation is marked by an arrow. fEPSP amplitude was normalized to baseline. (O) Magnitude of STP, determined as maximal responses within first 5 min after HFS, is significantly lower in DKO mice. Treatment of lithium further reduced the STP both in WT and DKO mice, but the significant interaction indicates a difference in dampening amplitude. (P) When normalized to mean STP in Ctrl treated mice, log2FC reduction of STP amplitude is significantly larger in wildtypes compared to DKO mice. (Q) Magnitude of LTP, determined as responses between 50 and 60 min after HFS, is significantly lower in DKO mice. Treatment of lithium further reduced the LTP both in WT and DKO mice, but the significant interaction indicates a difference in dampening amplitude. (R) When normalized to the mean LTP in Ctrl treated mice, log2FC reduction of LTP amplitude is significantly larger in wildtypes compared to DKO mice. * p < 0.05; ** p < 0.01; *** p < 0.001; n.s. not significant; p-values refer to univariate two-way ANOVA with Type 2 sum-of-squares; simple effects were tested in a similar but unifactorial ANOVA procedure; WT: wildtype mice; DKO: BHLHE40^-/-^/41^-/-^ mice; Ctrl: vehicle control; LiCl: lithium-treated; G: genotype main effect; T: treatment main effect; GxT: interaction effect; simple T: simple effects; DKO-T: DKO simple treatment effect; WT-T: WT simple treatment effect.

This sensor construct was transduced via AAV infection in WT and DKO cultured cortical neurons chronically pre-treated with lithium and vehicle. After baseline recording, we stimulated the culture with bicuculline, a competitive GABA_A_ receptor inhibitor, and quantified the activity peak (Figure 3B-C, Table S4). We found a strongly reduced excitability of DKO cortical neuron cultures and a dampening of the bicuculline response after lithium treatment only in WT neurons, whereas DKO neurons were non-responsive (GxT: P = 0.00196; WT-T: P = 0.00479; DKO-T: P = 0.375). To confirm that this reduced signal was caused by reduced neuronal network activity and not cytotoxic effects of lithium chloride, we silenced lithium-treated WT neuron cultures with a cocktail of TTX and AP-V, which fully eliminated the dampening effect (Figure S9A).

DKO mice display alterations in forebrain-dependent cognitive processing ^32^ and the lithium non-response was prominent in the cognitive domain, including learning flexibility, which is associated with the prefrontal cortex in rodents ^43^ and humans ^44^ and has been associated with BD^45^. Therefore, we compared the transcriptional profile of DKO and WT animals in the anterior cingulate cortex (ACC) of the medial prefrontal cortex (mPFC) at single-cell resolution to identify the most relevant cell type for further characterization (Figure S10). In order to find transcriptional signatures not modulated and dominated by daytime activity ^31^, we collected samples at zeitgeber time 4, the trough of *Bhlhe41* expression in rodents ^19,46^.

We observed clear transcriptional shifts in cell populations between DKO and WT upon UMAP clustering (Figure S10B – C). Based on marker gene expression, we could identify six major cell populations (Figure S10, Table S5). When comparing the transcriptional profiles of DKOs and WT in each cell type using differential expression analysis, we found substantial gene expression changes only in the EN L2/3 population (Table S5 and Figure S10D). In an enrichment analysis of gene ontology (GO) terms of the differentially expressed gene (DEG) set (Table S5 and Figure S10D), we found “regulation of postsynapse organization” to be the top enriched meta feature (−log(P) = 5.82, q = 0.033)(Figure S10F) and that the DEGs from L2/3 form a tight cluster with the top genes from the genetic analysis (Figure 1B). This suggests that the mPFC-associated phenotypes primarily depend on alterations of pyramidal cells in cortical layer 2/3, which receive monosynaptic input from ventral hippocampus (vHIP) via direct projections to the mPFC, in particular in the prelimbic (PL) and anterior cingulate cortex (ACC) (Liu and Carter 2018).

To elucidate lithium (non-) responsiveness in a more physiological context, we did whole-cell recordings on acute brain slices in two regions of interest that are associated with cognition: the ACC region of the mPFC and the CA1 region in the vHIP (Figure 3D). Mice were chronically treated for three weeks with LiCl or vehicle (Ctrl) via drinking water before the first recordings.

Based on our findings in single-cell transcriptomics (Figure S10D), we first focused on layer 2/3 pyramidal cells in the ACC. In these cells, we found that mEPSC and sEPSC amplitudes were dampened in lithium-treated WT and DKO pyramidal cells (Figure 3E, G). However, for miniature and spontaneous EPSCs, there was a statistically significant interaction of genotype and treatment (mEPSC: GxT: P = 0.0364; sEPSC: GxT: P = 0.0309), demonstrating a reduced lithium effect in DKO neurons. This was confirmed by normalizing to the respective vehicle control, showing a reduced dampening in DKO ACC slices compared to WT (Figure 3F, H; mEPSC: P = 0.0178).

In CA1 pyramidal cells, spontaneous but not mEPSC amplitudes were reduced in slices from lithium-treated mice (Figure 3I, K; mEPSC: T: P = 0.0822, GxT: P = 0.129; sEPSC: T: P = 9.82 × 10^−6^, GxT: P = 0.630) with no significant interaction effects. However, after normalization to the mean of vehicle-treated slices, we did find a significantly weaker dampening of the mEPSC amplitude in DKO neurons compared WT (Figure 3J; P = 0.00195), but no significant difference in sEPSC reduction (Figure 3L; P = 0.495). This also reflects a partial lithium non-responsiveness for the dampening effect on excitability, similar to what we have observed in cortical cultures.

No such effects were found for mIPSC and sIPSC amplitudes both in ACC and vHIP recordings (Figure S11A-H), indicating that the difference in bicuculline response found in cortical cultures was not due to changes in GABAergic activity.

For mEPSCs and sEPSCs of DKO Ctrl pyramidal neurons, amplitude and frequency were reduced compared to WT Ctrl (Figure S11I – L) and also the frequency of mIPSC and sIPSC was reduced despite their unchanged amplitude (Figure S11M-P).

Interestingly, lithium-treatment elevated mEPSCs and sEPSCs frequency in both ACC and vHIP of DKO pyramidal neurons and of mIPSC and sIPSC only in vHIP back to WT levels (Figure S11I – P).

To investigate changes in synaptic plasticity, we recorded long-term potentiation (LTP) in the CA1 region of the vHIP, which sends projections to the mPFC ^49^ (Figure 3M – Q). Here, we could see a reduced potentiation of fEPSP amplitude in DKO slices compared to WT as well as a dampening of similar magnitude in slices of WT lithium treated mice (Figure 3M). The input-output function did not reveal changes in basal synaptic transmission (Figure S12). We also found a highly significant interaction of genotype and treatment for both short-term (STP) and long-term potentiation (Figure 3N and P; STP: GxT: P = 1.36×10^-22^, LTP: GxT: P = 1.27 × 10^−5^). This is reflected by a significantly reduced dampening in response to LiCl in DKO slices compared to WT (Figure 3O and Q; STP: P = 3.27×10^-7^; LTP: P = 8.82 × 10^−17^).

This supports our assumption that a change in excitability is responsible for the lithium-(non-) response mediated by the clock modulators BHLHE40/41.

### Single-cell transcriptomics identifies alterations in the lithium-response most prominent in layer 2/3 cortical neurons

To dissect differential Li-responses in WT and DKO animals chronically treated with lithium, we performed transcriptomic profiling at single cell resolution again by sampling the ACC at ZT4 (Fig. 4A, Figure S10A).

**Figure 4.**
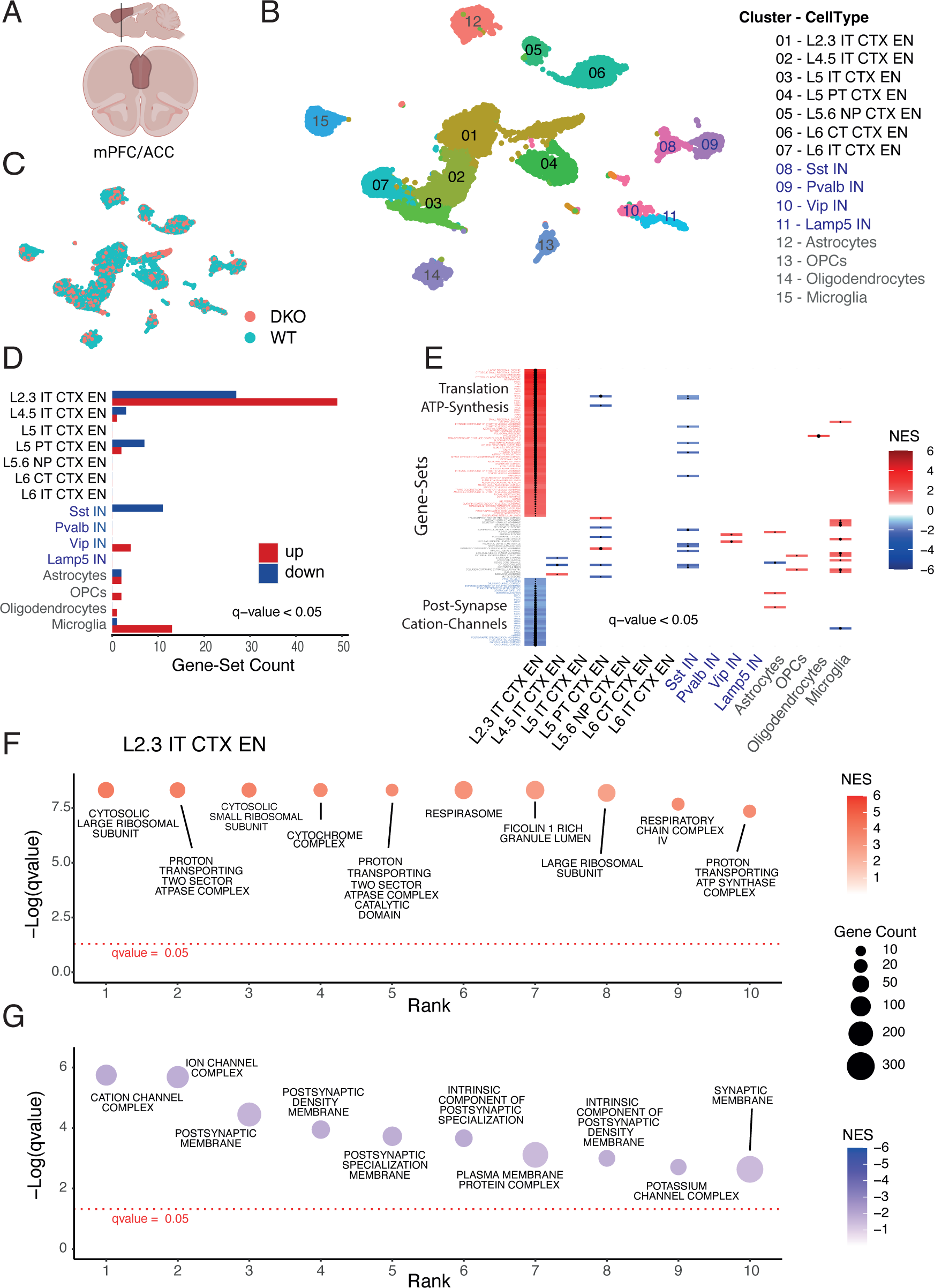
Lithium modulates transcription of ATP-synthesis, post-synapse, and cation-channel associated gene sets most prominently in Layer 2/3 excitatory neurons of the mPFC as assessed by RNAseq with single cell resolution. (A) The anterior cingulate cortex (ACC) portion of the medial prefrontal cortex (mPFC) was selected as region of interest. (B) UMAP dimension plot of lithium treated animals identifies 15 major cell populations, shown in distinct colors. Seven subclasses of excitatory cortical neurons, four subtypes of inhibitory neurons and four major glial cell types were identified (black, blue and grey coloured in the legend, respectively). (C) No differences in cell type clusters were detected between WT and DKO samples. (D) Barplot of significantly deregulated gene-sets (corrected q < 0.05) identified with GSEA against the gene ontology cellular component (GO-CC) collection shows most responses in layer 2/3 excitatory cells (49 up-and 27 down-regulated gene-sets) compared to all other cell types. (E) Condensed heatmap of all up-and down-regulated gene-sets (corrected q < 0.05) ordered for q-values shows the highly layer 2/3 specific response architecture with only sparsely overlapping gene sets enriched in other cell types. (F,G) Dotplot of the ten most significantly up-(F) and down-regulated (G) gene-sets ranked by the-log10 transformed q-value showing 6/10 upregulated gene-sets associated with mitochondrial ATP-synthesis and 3/10 with ribosome associated protein translation (F) and 5/10 and 3/10 down-regulated gene-sets associated with the post-synapse and cation-channel activities. L: cortical layer, EN: excitatory neurons, IT: intratelencephalic, PT: pyramidal tract, NP: near-projecting, CT: corticothalamic, IN: inhibitory neurons; Sst: Somatostatin, Pval: Parvalbumin, Vip: Vaso-Intestinal Peptide, OPC: oligodendrocyte precursor cell, NES: nominal enrichment score.

Based on marker gene expression, we identified 15 cell populations in both genotypes (Figure S13, Figure 4B, C, Table S6). Differential gene expression analysis revealed that substantial changes were again most prominent in excitatory neurons of cortical layers 2/3 in comparison to all other neuronal and glial cell types (Figure S14A). When comparing the transcriptional profiles upon lithium treatment in each cell type using differential expression analysis at the individual gene level, we found several genes to be downregulated in layer 2/3 excitatory neurons in lithium treated DKOs compared to WT which are associated with synaptogenesis (Lrrtm4, Nrg1), synaptic functions (Dlg2, Nlgn1, Nlgn2, Nrxn3), and cation channel function and excitability (Kcnip4, Cacnb2) (Figure S14B and Table S6). Among the upregulated genes were a subunit of the Na^+^/K^+^-ATPase (ATP1A3) a metabolic enzyme (Aldoa) and a ribosomal subunit (Cmss1) (Figure S14B and Table S6).

Next, we performed a more sensitive analysis on gene-set level (GSEA)^50^ by probing the transcriptomes of all 15 cell types against four well-annotated gene-ontology and pathway collections (Reactome and the Gene Ontology subcollections ‘Cellular Component’ (GO-CC), ‘Biological Porcess’ (GO-BP), and ‘Molecular Function’ (GO-MF). This analysis revealed that most significantly deregulated gene-sets were detected in layer 2/3 excitatory neurons (Figure 4D, Figure S15, Table S7) matching the analysis at the gene-level (Figure S14A). Plotting all gene-sets identified in all cell types (with a stringent q-val<0.05 cutoff) ranked by significance in a heatmap clearly showed the non-overlapping and highly selective layer 2/3 neuron specific response architecture (Figure 4E). The most prominent upregulated gene-sets were associated with ribosomal function/translation and mitochondrial respiration/ATP-synthesis (Figure 4E, F, Figure S15, Table S7), whereas cation-channel and post-synapse associated gene-sets were among the top downregulated pathways (Figure 4E, G, Figure S15, Table S7). We dissected the synapse-associated enrichments further by a focused SynGO localization and function query ^51^. Cellular component categories showed enrichment particularly but not exclusively for the ‘postsynapse’ in untreated and lithium treated samples (corrected p-val: untreated = 3.6 x 10^-6^; lithium = 4.4 x 10^-9^) and ‘postsynaptic density’ (corrected p-val: untreated = 1.0 x 10^-4^; lithium = 2.1 x 10^-7^) (Table S8, Figure 5A). Functionally, the central category ‘process in the synapse’ was also highly significantly enriched in untreated and lithium treated samples (corrected p-val: untreated = 6.0 x 10^-7^; lithium = 4.5 x 10^-11^). Enrichment of genes involved in ‘synapse organization’ was dramatically increased upon lithium treatment (corrected p-val: untreated = 0.01; lithium = 6.8 x 10^-11^) (Table S8, Figure S17). We generated an extended protein network of all DGEs that formed a tight cluster of almost all genes (Figure 5B). Highlighting synaptic genes corresponding to the above mentioned SynGO categories showed the centrality of synaptic mechanisms underlying the lithium response architecture (Figure 5C-E).

**Figure 5.**
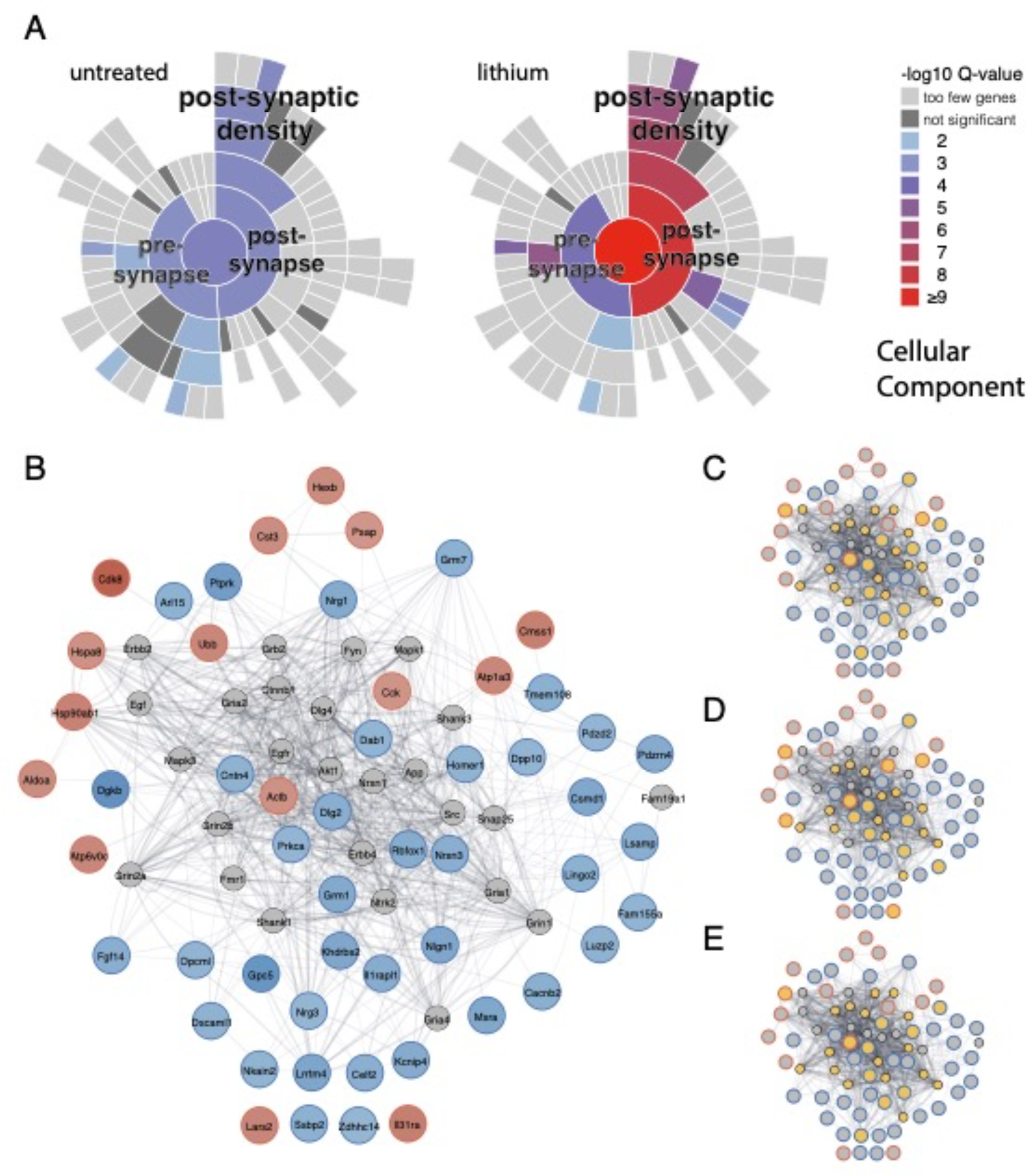
Enrichment of Lithium modulated genes in synaptic structures in Layer 2/3 excitatory neurons from wildtype and DKOs. (A) SynGO plot of enrichment of gene sets associated with synaptic structures in wt (left) and DKOs (right). Significantly associated genes are found in the top-layered pre-and postsynapse and branch our most prominently into post-synaptic membrane associated structures. (B) Protein network of lithium deregulated genes forms a highly interconnected cluster (Up-and downregulated gene products are depicted as blue and red circled nodes, respectively. Protein-protein-interactions (PPI) are plotted as greyish lines between nodes. Nodes added by the PPI network to increase connectivity are plotted as smaller sized grey circles. Only 4 genes are not integrated into the main network based on published PPIs (C-E) Synaptic components from the SynGO pre-synapse (C), post-synapse (D) and post-synaptic density (E) subclusters are highlighted in yellow and are enriched at the ‘synaptic core’ of the network.

Taken together, single cell RNAseq analyses identified a highly specific cortical layer 2/3 excitatory neuron response modulated by lithium and BHLHE40/41. The most prominent deregulated genes and gene sets are associated with mitochondrial respiration, translational control and synaptic functions.

## Discussion

In this study, we have applied gene set enrichment methods to identify human clock genes relevant to lithium therapy outcome in BD patients. The strong association with chronotype as a positive control trait, the validation with two independent methods (MAGMA and INRICH ^29,30^) and two independent patient cohorts, as well as the translational and functional validation in two transgenic mouse models proved the robustness of our approach. Our results confirmed a global influence of the core clock gene set on the lithium responsiveness but did not corroborate an association with BD as a trait ^52^. Thus, this specific association at the gene set level further supports the assumption that the genetic architecture of lithium treatment response contains biological components largely independent from other psychiatric traits. Although some core clock genes seem to be associated selectively with lithium responsiveness (*CSNK1E*, *BHLHE41*), others show a mixed (*RORB*) or selective association with BD (*ARNTL*), arguing for independent and overlapping neurobiological mechanisms modulated by clock genes between lithium response and BD disease root causes.

CSNK1E is a kinase that modulates many circadian processes and has been shown to directly phosphorylate PER1 and PER2 (Maywood et al. 2014). Mouse loss of function mutants display deficits in affective behaviors and sleep architecture ^55,56^. Although many studies have provided indirect evidence for an involvement of CSNK1E in BD ^57^, our GWAS analysis associated the gene locus with lithium-response and not BD life-time risk as a trait. Therefore, future studies are needed to functionally link this gene with lithium-response. *BHLHE41* (also known as *SHARP1* or *DEC2*) codes for a transcriptional repressor that regulates the activity of clock and clock-controlled genes. In our study, we found an association with lithium-response but not BD risk. Mutations in this gene reduce sleep length in humans and rodents ^58,59^ and *Bhlhe40/41*^−/−^ double-knockout (DKO) mice are characterized by altered sleep patterns, cognitive alterations and mixed-state behavioral endophenotypes of psychiatric disorders ^19,32^. RORB (RAR Related Orphan Receptor B) is a member of the NR1 subfamily of nuclear hormone receptors. It binds as a monomer to ROR response elements and plays a key role in the regulation of the period length and stability of the circadian rhythm ^60^. Our analysis associated RORB with lithium-response, but the genetic association with BD life-time risk was more pronounced. *ARNTL* (also known as *BMAL1* and BHLHE5) is the only core clock gene that is indispensable for the maintenance of the circadian cycle ^60^. At the gene level, we did not detect an association with lithium-responsiveness but could validate its association with BD risk.

We focused on BHLHE41 for further analyses, because of its specific association with lithium response. We performed a comprehensive behavioral, molecular, and physiological characterization of the lithium-response in BHLHE41/41 double null mutant mice (DKOs), since both genes are functionally redundant with respect to circadian and cognitive phenotypes ^31,32^. Although lithium had a strong impact on arousal-related measures and 24h activity profiles in WT and DKOs, highly significant genotype-treatment interactions were detected mainly in the cognitive domain. In contrast to *Arntl* Het mice, DKOs were not responding to chronic lithium treatment like wildtype mice, most evident in the recent and remote contextual fear memory tasks. This finding argues for hippocampus and cortex dependent mechanisms underlying the lithium non-responsiveness. Thus, we focused our molecular and electrophysiological analyses on the cortex and hippocampus, and furthermore because of the absence of lithium dependent alterations of circadian activity measures.

The ACC and hippocampus are known to be involved in BD pathology, e.g. BD patients have reduced ACC grey matter volume (Ellison-Wright and Bullmore 2010) and reduced hippocampal volume ^63^ and the ACC is implicated in the control cognition and emotional states in BD^45^. Interestingly, hippocampus and cortical grey matter thickness were increased in lithium-treated patients compared to other BD patients ^64,65^. As mentioned above, the neuroprotective effect was mostly shown *in vitro*, e.g. the non-responsiveness of long-term lithium-treated excitatory neuron cultures to excitotoxicity ^1,66^. These findings are in line with the reduced ESARE signal that we found in response to stimulation in primary cortical neurons. It indicates an already lowered level of excitability in DKO cortical neurons that was not reduced further by lithium treatment. We could validate the dampened neuronal excitability in acute brain slices, finding smaller mEPSC and sEPSC amplitudes in layer 2/3 ACC and CA1 pyramidal cells in DKO slices as well as a reduced dampening effect of chronic lithium treatment. Reduced excitability has also been detected in patient derived human neurons and cortical spheroids derived from Li-non-responders ^11,13^.

Interestingly, mEPSC and sEPSC frequencies were also lower in control DKO slices compared to WT but came back to WT levels in lithium treated samples. These findings suggest that the response of DKO pyramidal cells to lithium was not only reduced, but fundamentally different from WT pyramidal cells, likely resulting in a difference in synaptic plasticity and producing network level changes different from WT. In support of these observations, BHLHE41 and BHLHE40 were associated with daytime neuronal activity regulation in the past ^31,46^. The transcription of BHLHE40 was also found to be induced in response to kainate in rat cerebral cortex ^38^ and a knockdown dampened synapse-to-nucleus signaling in a previously published ESARE assay ^41^. Furthermore, DKO mice were found to exhibit a dampened circadian immediate early gene (IEG) expression amplitude in the cortex ^19^.

Transcriptomic profiling at cellular resolution with ACC samples harvested from untreated and lithium treated DKO and WT animals revealed a highly cell type specific response restricted to excitatory neurons of cortical layers 2/3. The most prominent gene sets found upregulated in DKO layer 2/3 neurons versus WT baseline were associated with cellular cation homeostasis and the organization of the post-synapse. In addition, and specifically detected only upon lithium treatment, gene sets associated with mitochondrial respiration and ATP synthesis as well as ribosomal functions were upregulated in layer 2/3 neurons from DKOs compared to wildtype. Similar response profiles at the pathway level have been obtained with human iPSC derived neural precursor cells and brain organoids ^12,13^. Together, these results imply that chronic lithium treatment induces responses at multiple levels likely involving different cation-associated mechanisms causing changes in membrane excitability and mitochondrial proton gradients. The response of synaptic and ribosomal gene expression profiles might be an indirect adaptation of the above-mentioned interference of lithium with the different cation-controlled mechanisms operating in neurons that link metabolic states with synaptic functions. The astonishingly cell type specific response to lithium treatment that we see in the adult cortex of mice may be explained by differences in cellular uptake and/or subcellular retention rates mediated by differential expression of channels and/or transporter proteins ^13,67^. Brain region dependent lithium uptake and distribution of candidate channels/transporters has been described ^68^, our results show that further analyses at the cell type level will be critical to better understand lithium’s’ mode-of-action in living animals and patients.

In this study, we have identified components of the molecular clock including BHLHE40/41 as critical modulators of the lithium response in BD patients. We show that lithium treatment lowered the excitability of WT cortical neurons, whereas BHLHE40/41 mutant neurons were less responsive. Moreover, we found that BHLHE40/41 mutant pyramidal cells are less responsive to lithium-induced reduction in mEPSC and sEPSC amplitude. Nearly complete lithium non-responsiveness was seen in LTP of the CA1 region of the hippocampus, and a non-responsiveness of DKO mice to lithium’s effect on vHIP-mPFC-dependent learning and memory. Thus, BHLHE40/41 mutants represent a promising mouse model to test novel treatments with the aim overcome lithium non-responsiveness in the future.

## Acknowledgements

We like to acknowledge the contribution of the whole team of the animal facility and behavioral unit of the Rossner lab for their continuous efforts. We also like to thank Meino Rohlfs (LMU University Childrens’ Hospital) for access and support with Illumina sequencing. Funding received: Department of Veterans Affairs Grant number BX003431 (MMC). Bundesministerium für Bildung und Forschung Grant number 01EK2101D (MJR). ECS was supported by the Munich Clinician Scientist Program (MCSP). The following conflict of interest statements are declared: MMC has served in 2022 and 2023 on the scientific advisory board for the Alkermes Pharmaceuticals Pathways Young Investigator Award. NK is employee and MJR shareholder of Systasy Bioscience GmbH. None of the companies has contributed to the design and analyses of the study.

## DISPLAY ITEMS

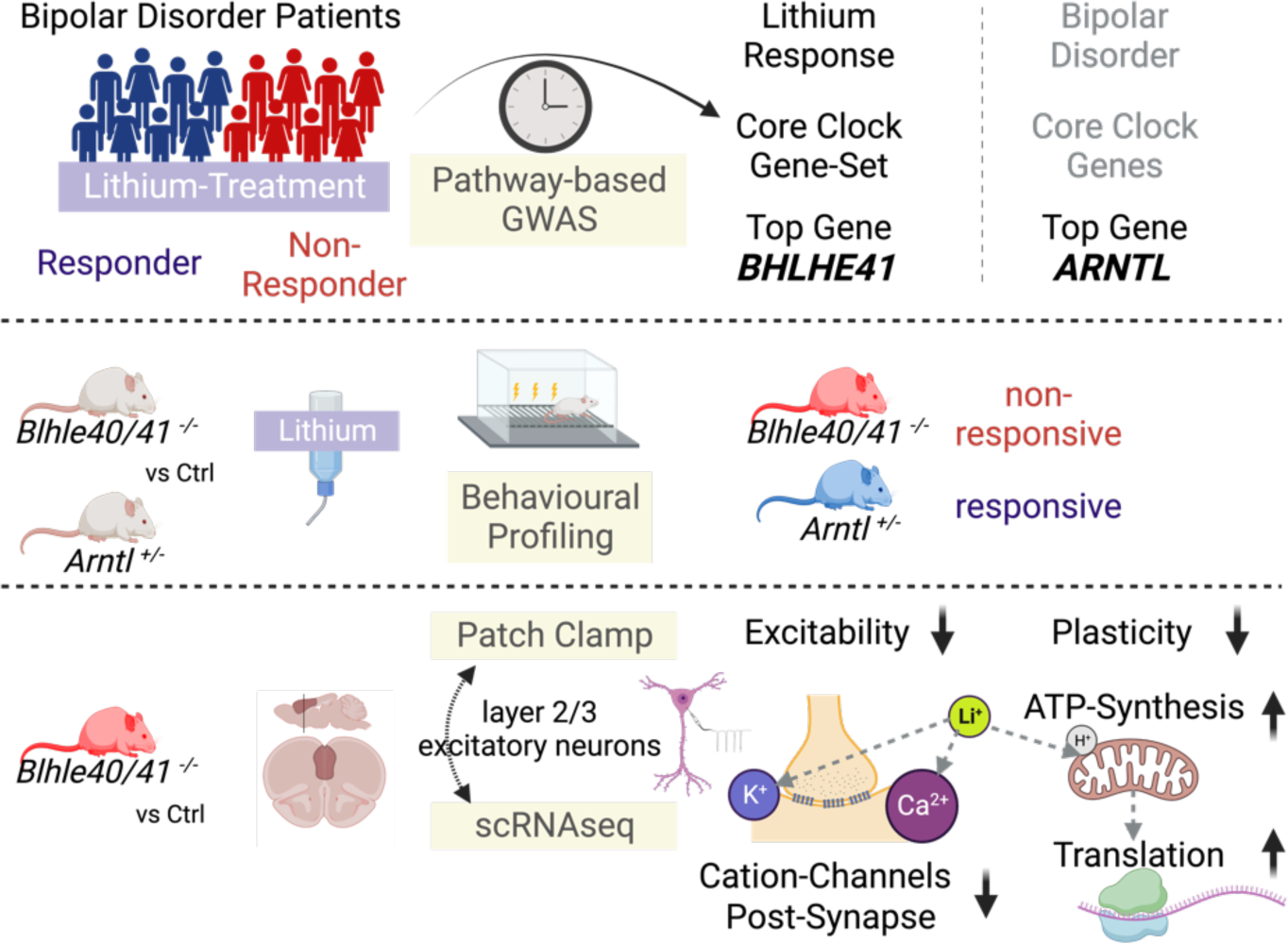
Grafical Abstract.

## Supplementary Figures

**Figure S1.**
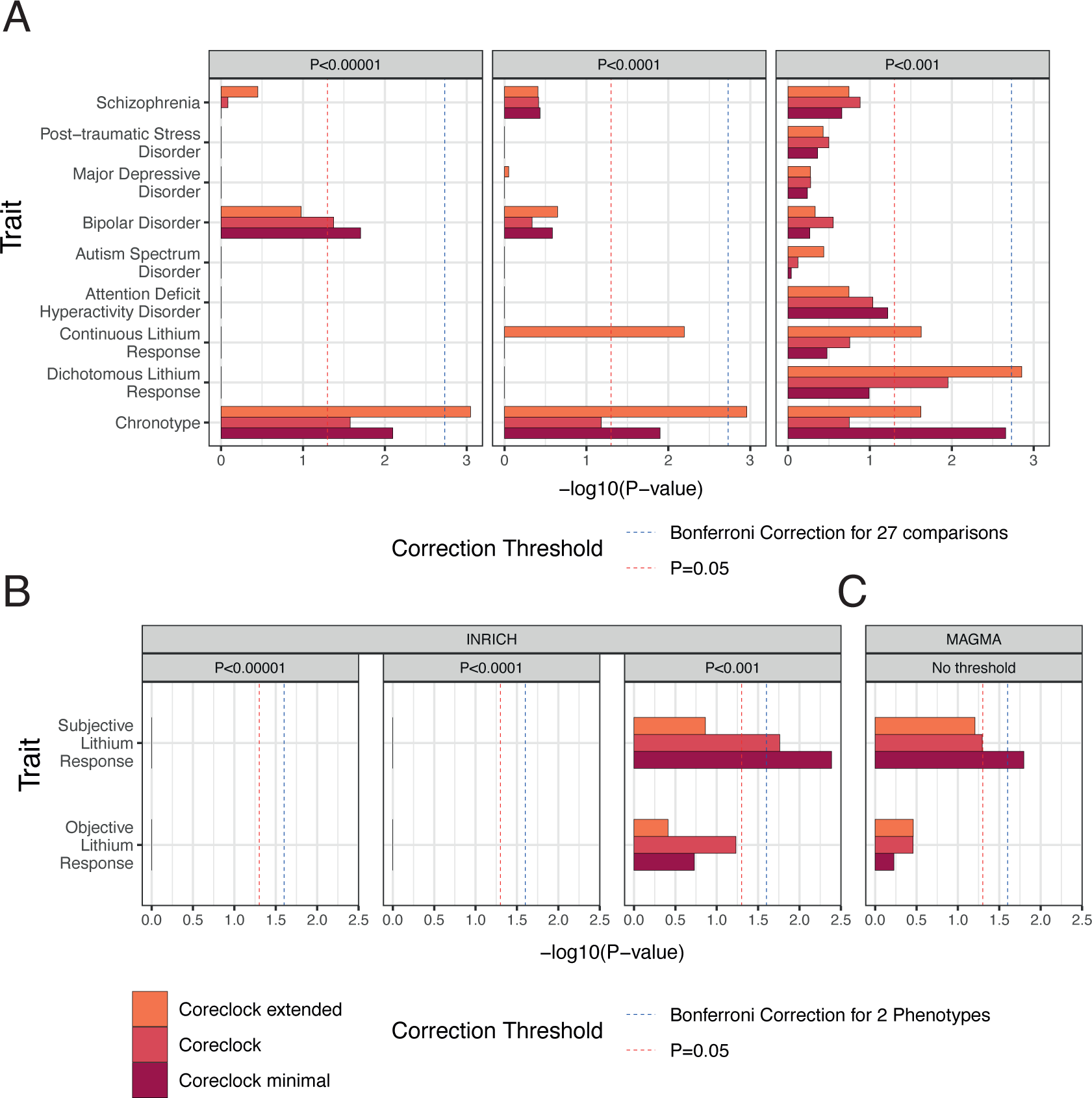
INRICH analyses based on core clock gene-sets in psychiatric traits. (A) INRICH analyses of data shown in Figure 1. Bonferroni significance thresholds: dashed blue line (P < 0.05/ (3 gene-sets x 9 traits)). Three different P-value thresholds were analyzed (P=0.00001, P=0.0001, P=0.001). The ‘Coreclock extended’ gene-set reaches corrected p-val-threshold at P=0.00001 and P=0.0001 in the Chronotype sample, whereas it reaches corrected p-val-threshold in the ‘Dichotomous Lithium Response’ sample at P=0.001. (B) INRICH analyses and (C) MAGMA enrichment analyses of swedish replication cohort with two phenotypes: subjective and objective lithium response. Bonferroni significance thresholds: dashed blue line (P < 0.05/ (3 gene-sets x 2 traits)). (Competitive test P-values in the X-axis are –log10 converted. Dashed red line indicates the nominal (uncorrected) significance threshold P=0.05.

**Figure S2.**
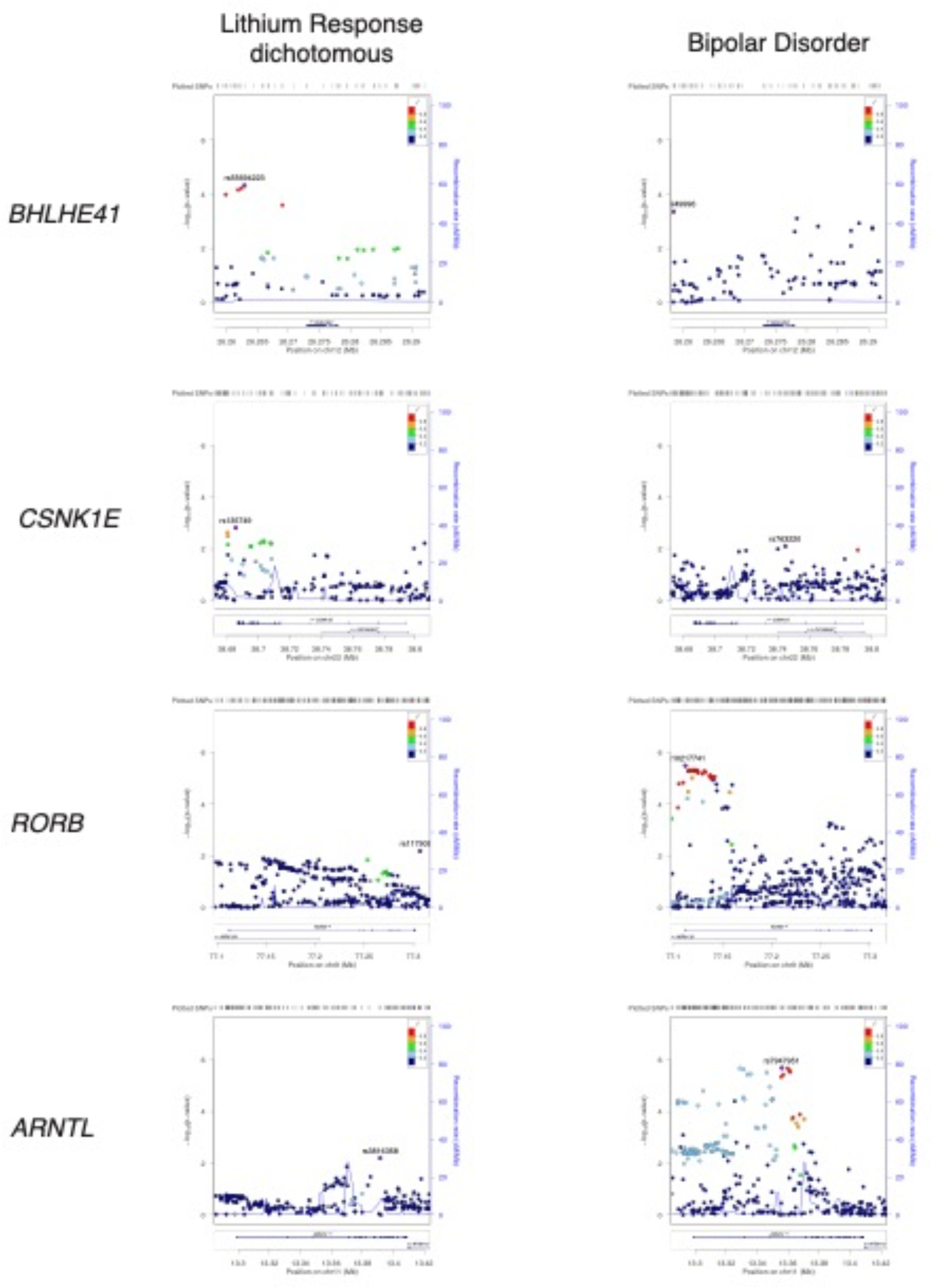
Manhattan plots for nominally significant contributing genes in the core clock gene-set and ARNTL. Manhattan plots from GWAS data on dichotomous lithium response (A, C, E, G) and bipolar disorder risk (B, D, F, H). For the top three (and nominally significant) lithium response-associated genes from the core clock gene-set: (A, B) *BHLHE41*, (C, D) *CSNK1E*, and (E, F) *RORB*. In addition, the same diagrams are shown for *ARNTL* (G, H), a GWAS associated BD risk gene also known as the core clock regulator *BMAL1*.

**Figure S3.**
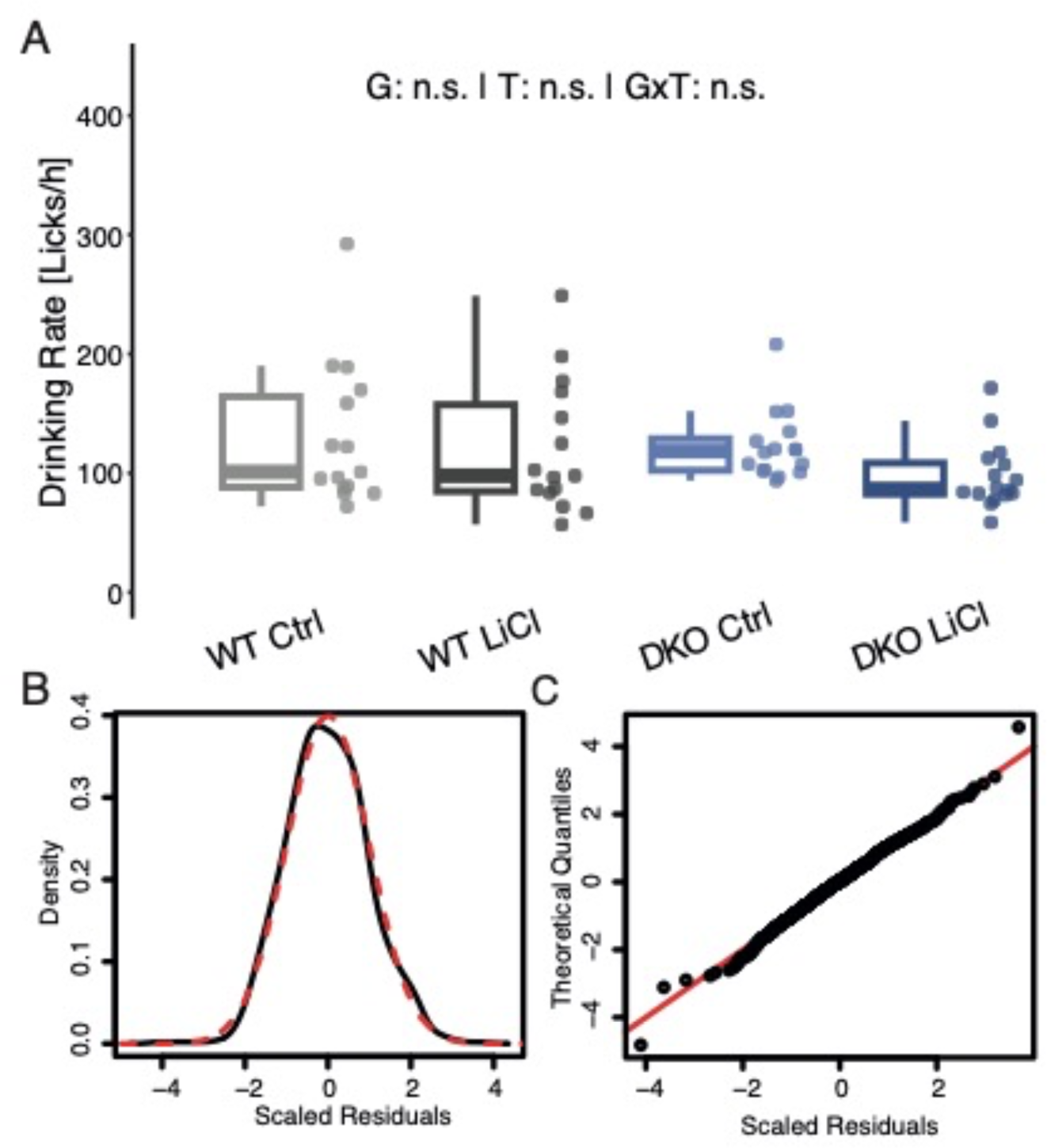
Quality controls for the PsyCoP behavioral study. (A) Drinking rate in the IntelliCage during the experiments was not significantly different in lithium chloride-treated mice compared to control. Data are shown as box plots with whiskers extending to no more than 1.5-fold IQR. (B) Density plot and QQ plot of the scaled residuals (black) of the multivariate linear model used for the ANOVA procedure of all PsyCoP measures compared to a normal distribution (red) show no major deviations. * P < 0.05, ** P < 0.01, *** P < 0.001, n.s. not significant; P-values are FDR-adjusted and refer to Wilk’s lambda testing two-way ANOVA; G: genotype term; T: lithium treatment term; GxT: interaction term; WT: wildtype; DKO: Bhlhe40/41 double-knockout; Ctrl: vehicle control; LiCl: lithium-treated

**Figure S4.**
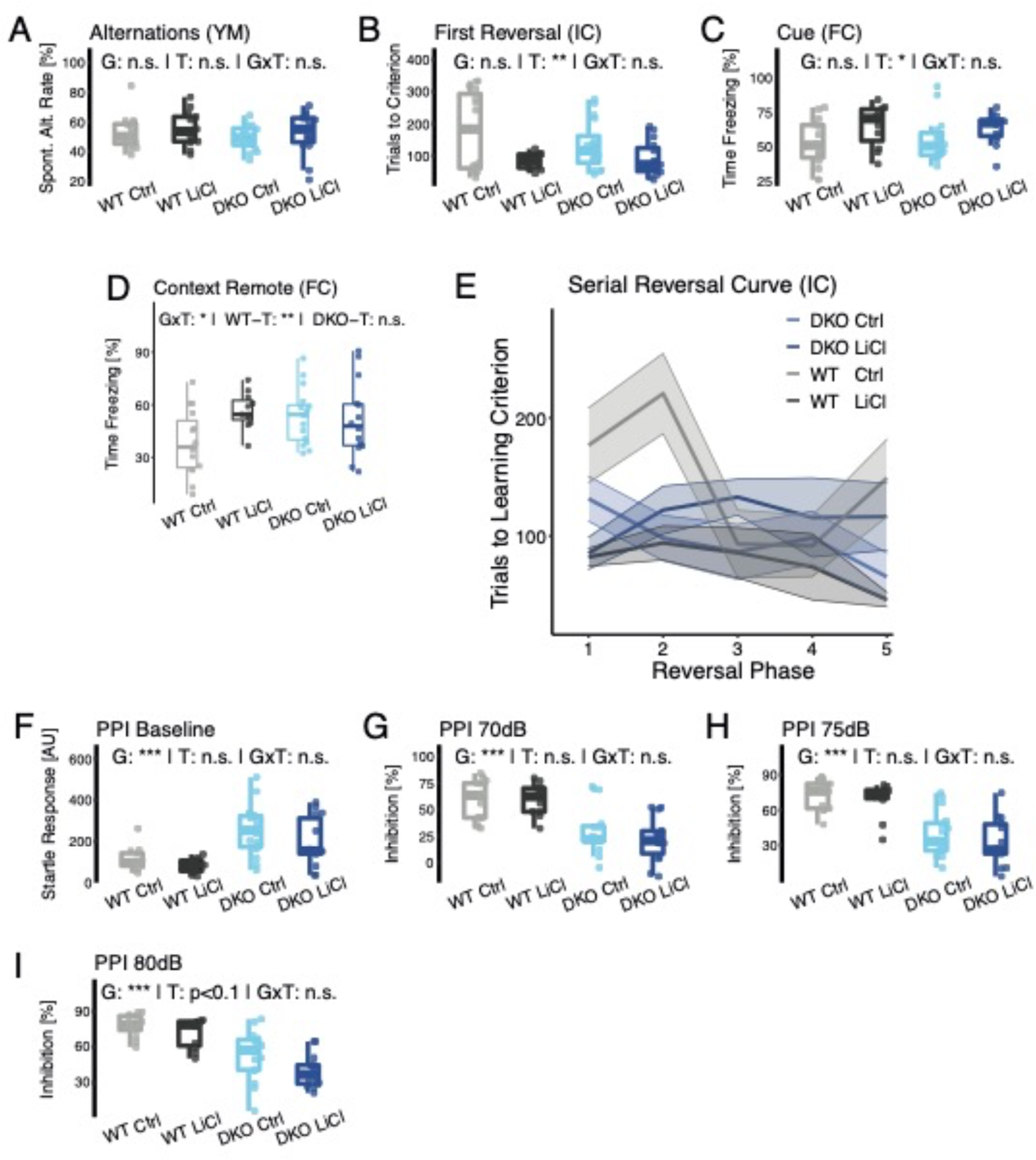
Remaining variables of the cognition and sensorimotor RDoC domains. (A-C) display the variables of the Cognitive RDoC domain not shown in Figure 2. (A) There was no significant effect on working memory in the Y-Maze test measured with the rate of spontaneous alternations between arms. (B) In the first reversal lithium treated mice needed less trials to reach the learning criterion indicating improved learning flexibility. Although, the difference in DKO mice was smaller than in wildtypes, the interaction was not significant in this part of the spatial learning experiments. (C) Similarly, in the cued fear memory task, lithium treatment increased the time spent freezing, showing improved cued fear memory, with no significant influence in DKOs. (D) In the remote contextual fear memory, however, there was a significant Li-treatment dependent effect between the genotypes, as indicated. (E) The ribbon plot of this curve refers to the Serial Reversal Task analysis (Fig. 2E) and displays the mean number of trials each group needed to reach the spatial learning criterion (10% better success rate than random expectation) after each reversal of the correct corner. Smaller values equal less trials needed and indicate faster learning success. WT animals improve upon Lithium treatment, whereas DKO performance remains unaltered. (F-I) show the variables of the Sensorimotor domain with a strong genotype effect in all of them, but only minor influence of lithium treatment on the highest prepulse level (PPI 80dB) in panel (I). Data are shown as box plots with whiskers extending to no more than 1.5-fold IQR; * P < 0.05, ** P < 0.01, *** P < 0.001, n.s. not significant; P-values are FDR-adjusted and refer to Wilk’s lambda testing two-way ANOVA; G: genotype term; T: lithium treatment term; GxT: interaction term; WT: wildtype; DKO: Bhlhe40/41 double-knockout; Ctrl: Placebo control; LiCl: lithium-treated

**Figure S5.**
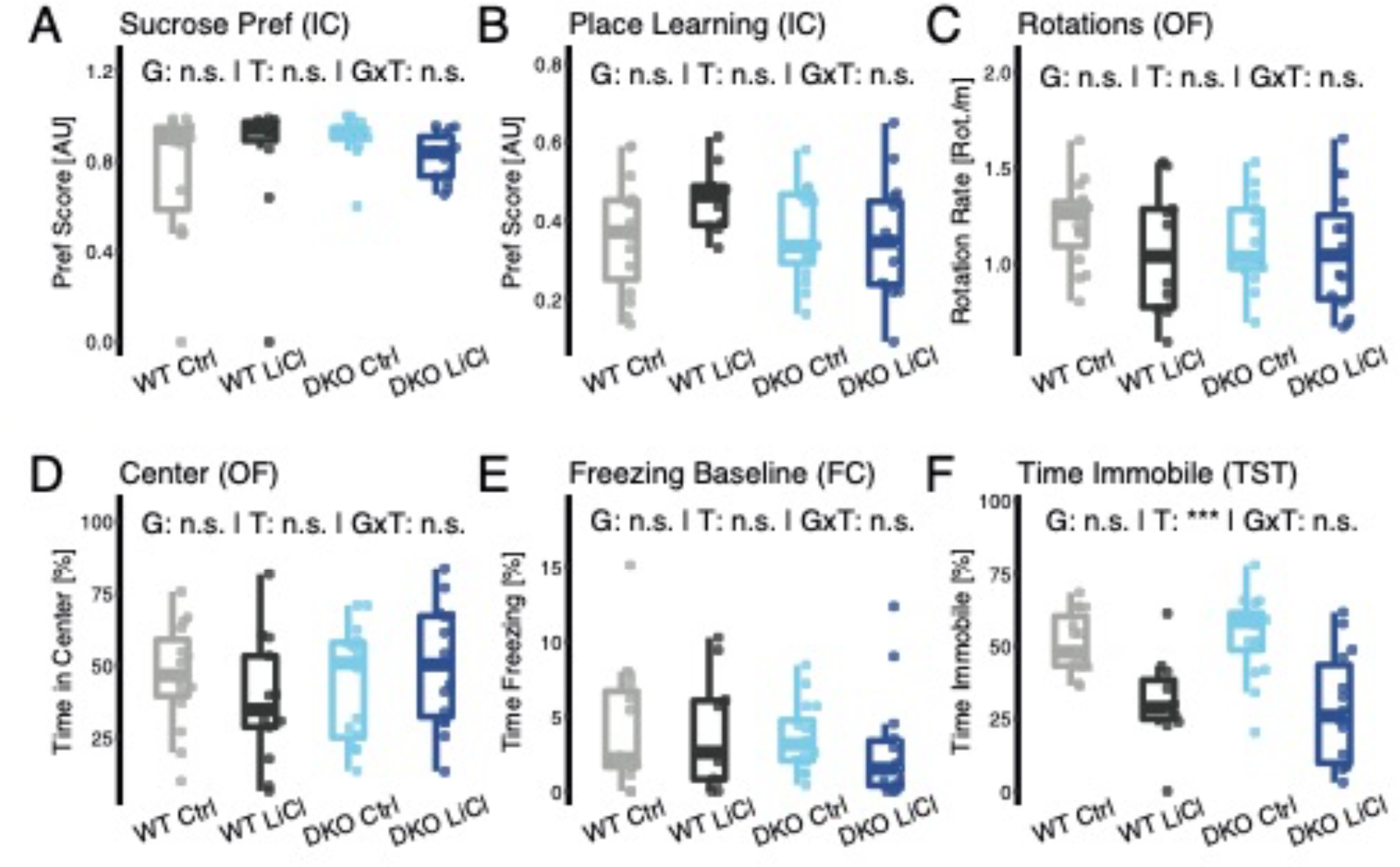
Positive and Negative Valence RDoC domains. The variables associated to positive valence systems did not show significant effects, neither in the Sucrose Preference Test (A) nor the place preference in Positive Reinforcement Learning (B). Notably, there was sex difference in sucrose preference detected, shown in Figure S7. In the negative valence domain parameters, there were no significant differences in rotation rate (C) and the center time (D) in the Open Field Test as well as the baseline freezing behavior (E) in Fear Conditioning either. However, lithium-treated mice were significantly less immobile (F) in the Tail Suspension Test independent of genotype. Data are shown as box plots with whiskers extending to no more than 1.5-fold IQR; * P < 0.05, ** P < 0.01, *** P < 0.001, n.s. not significant; P-values are FDR-adjusted and refer to Wilk’s lambda testing two-way ANOVA; G: genotype term; T: lithium treatment term; GxT: interaction term; WT: wildtype; DKO: Bhlhe40/41 double-knockout; Ctrl: Placebo control; LiCl: lithium-treated

**Figure S6.**
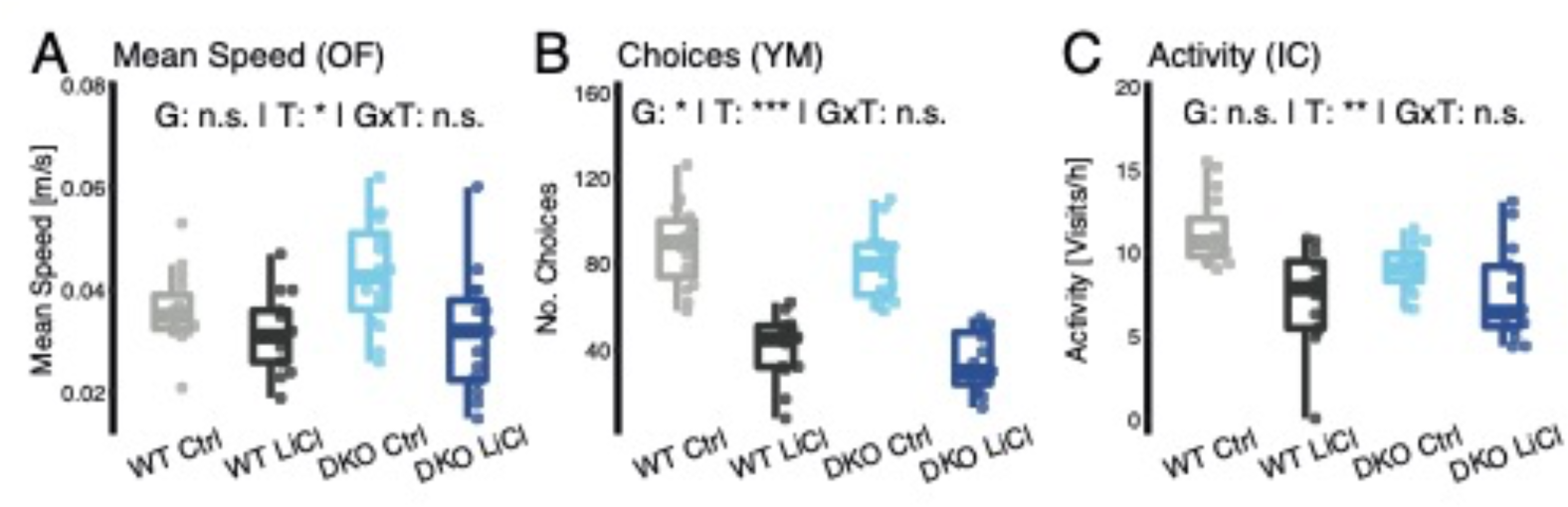
Arousal and Regulatory RDoC domain. Novelty-induced activity was reduced in lithium-treated animals (A) measured as Mean Speed in the Open Field Test and (B) the number of arm choices in the Y-Maze Test. In latter case Bhlhe40/41 DKO mice displayed a slightly reduced activity, as well, independent of treatment. General Activity (C) in the IntelliCage was similarly reduced in response to lithium treatment. Please note that there was a sex difference in activity level in response to lithium treatment shown in Figure S7 with males being impacted stronger than females. Data are shown as box plots with whiskers extending to no more than 1.5-fold IQR; * P < 0.05, ** P < 0.01, *** P < 0.001, n.s. not significant; P-values are FDR-adjusted and refer to Wilk’s lambda testing two-way ANOVA. G: genotype term; T: lithium treatment term; GxT: interaction term; WT: wildtype; DKO: Bhlhe40/41 double-knockout; Ctrl: Placebo control; LiCl: lithium-treated

**Figure S7.**
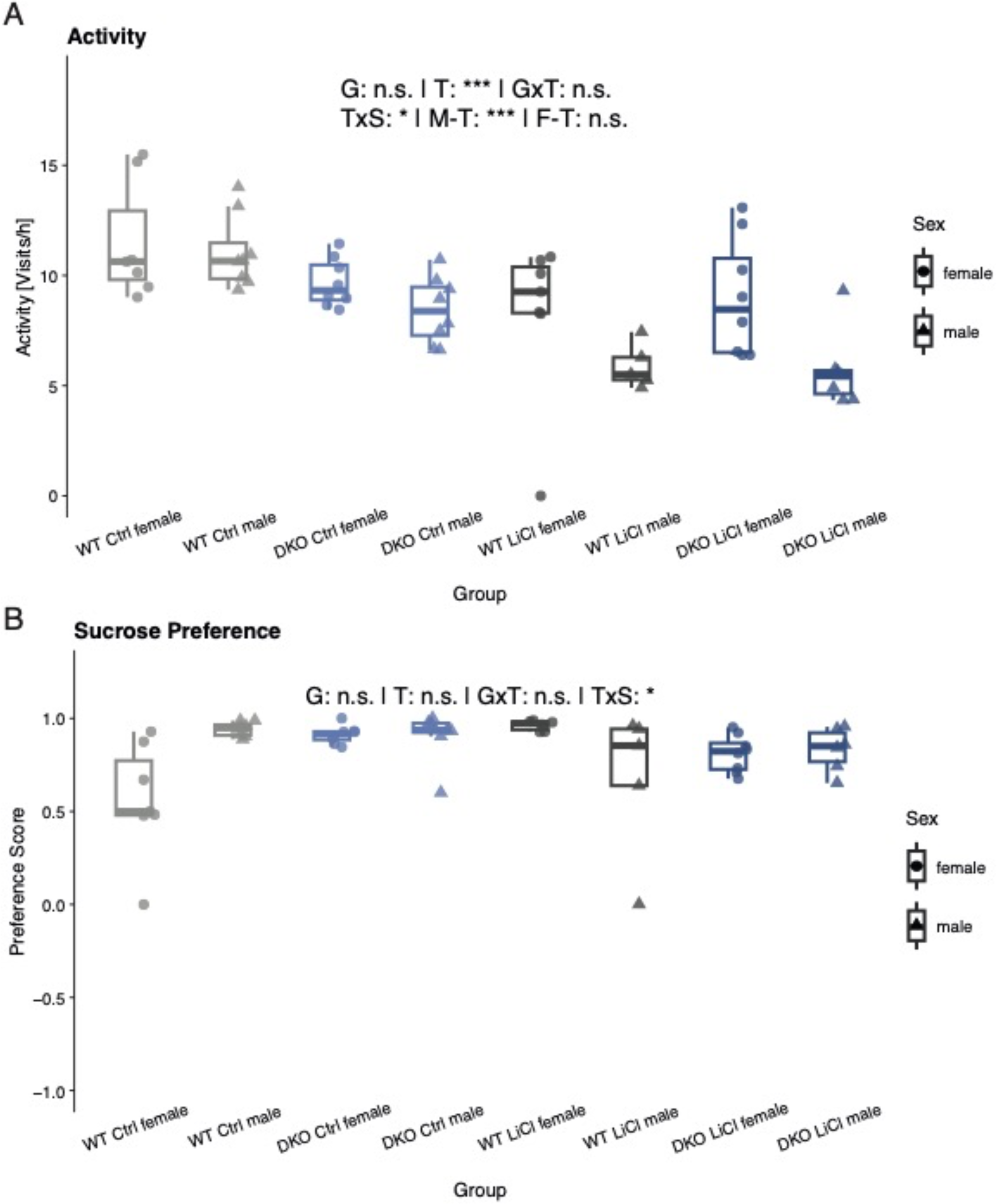
Sex Differences in general activity and sucrose preference. In most behavioral parameters assessed, we did not detect significant differences between sexes (Table S3). (A) However, male mice showed a stronger reduction in general activity in response to lithium treatment than female mice. (B) Moreover, the female wildtype placebo control (WT Ctrl female) group displayed a lower sucrose preference than their lithium-treated counterpart. This difference in lithium response between sexes was not found in DKO mice. Data are shown as box plots with whiskers extending to no more than 1.5-fold IQR; * P < 0.05, ** P < 0.01, *** P < 0.001, n.s. not significant; P-values are FDR-adjusted and refer to Wilk’s lambda testing two-way ANOVA; n =; G: genotype term; T: lithium treatment term; GxT: G by T interaction term; TxS: T by sex interaction term; M-T: male treatment effect; F-T: female treatment effect; WT: wildtype; DKO: Bhlhe40/41 double-knockout; Ctrl: Placebo control; LiCl: lithium-treated

**Figure S8.**
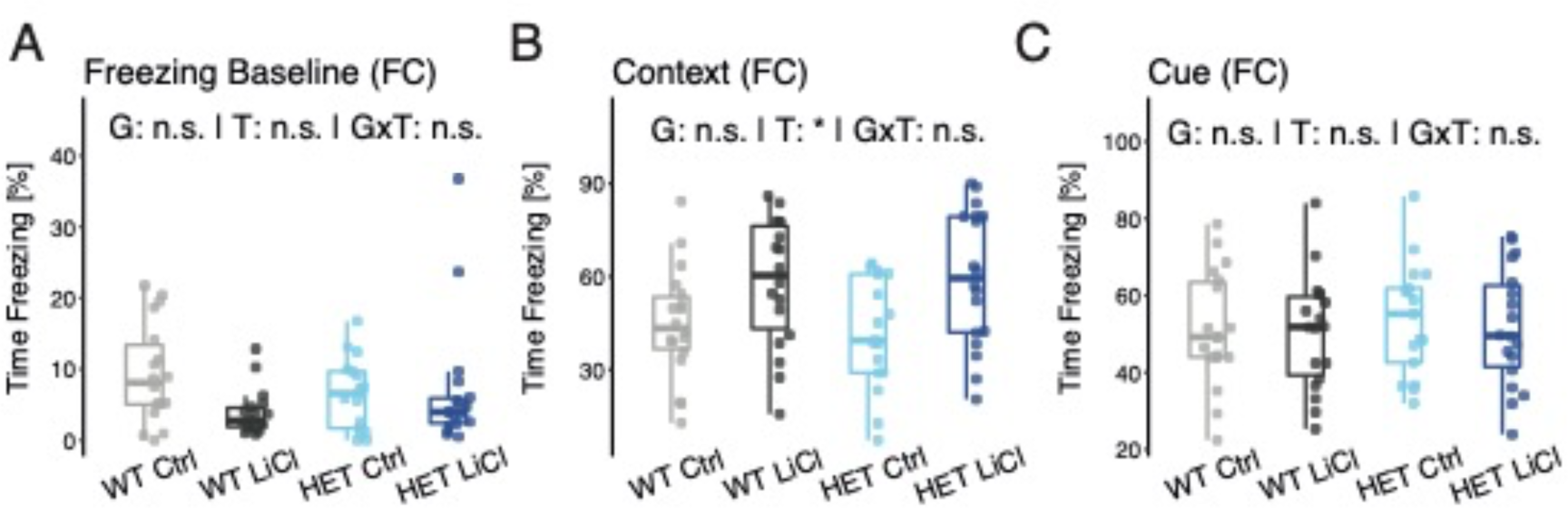
Fear Conditioning of Arntl mouse model. (A-C) Arntl heterozygous null mutant mice (Het) and littermate controls untreated or treated with lithium were subjected to fear conditioning (FC) paradigm. (A) No differences were detected at baseline freezing. (B) Testing contextual fear revealed a treatment dependent significant increase in both genotypes. (C) Cued FC detected no changes. Data are shown as box plots with whiskers extending to no more than 1.5-fold IQR; * P < 0.05, ** P < 0.01, *** P < 0.001, n.s. not significant; P-values are FDR-adjusted and refer to Wilk’s lambda testing two-way ANOVA; n =; G: genotype term; T: lithium treatment term; GxT: G by T interaction term; TxS: T by sex interaction term; M-T: male treatment effect; F-T: female treatment effect; WT: wildtype; Het: Arntl heterozygous null mutants; Ctrl: Placebo control; LiCl: lithium-treated

**Figure S9.**
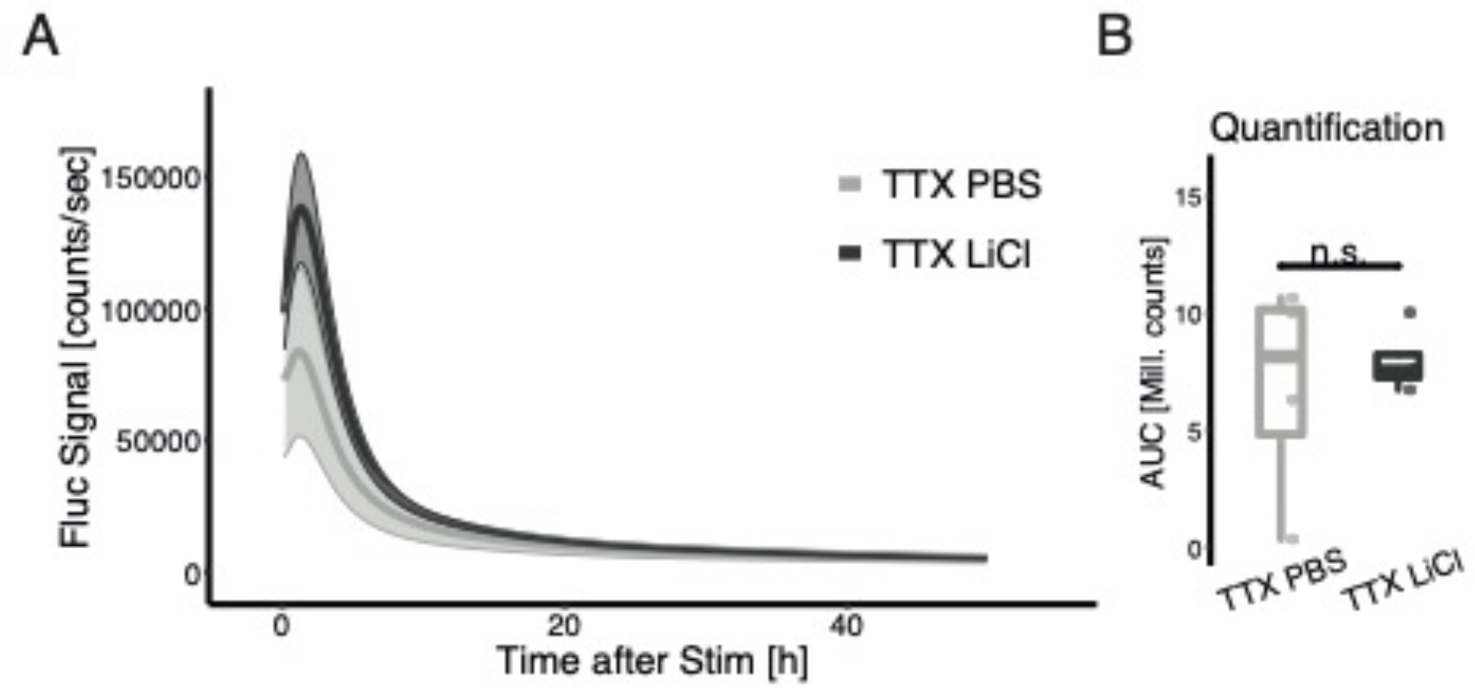
Lithium toxicity. (A) First 50 hours of tetrodotoxin (TTX)-and AP-V-silenced primary cortical neuron culture from wildtype (WT) embryos after lithium treatment. Cells were transduced with an ESARE-Firefly Luciferase (Fluc) reporter construct via AAV infection. After 7 days of pretreatment with lithium chloride (LiCl), a TTX/AP-V cocktail was added to silence neuronal network activity. At the start of the recording a typical artifact peak resulting from the time-lag of the transcriptional-translational reporter system, was seen. (B) The area under the curve of the first 100 hours of recording was quantified. No statistically significant difference was found in a Student‘s two-sided t-test, indicating that differences in ESARE-response are due to activity dampening not cytotoxic effects of lithium chloride. n.s.: not significant.

**Figure S10.**
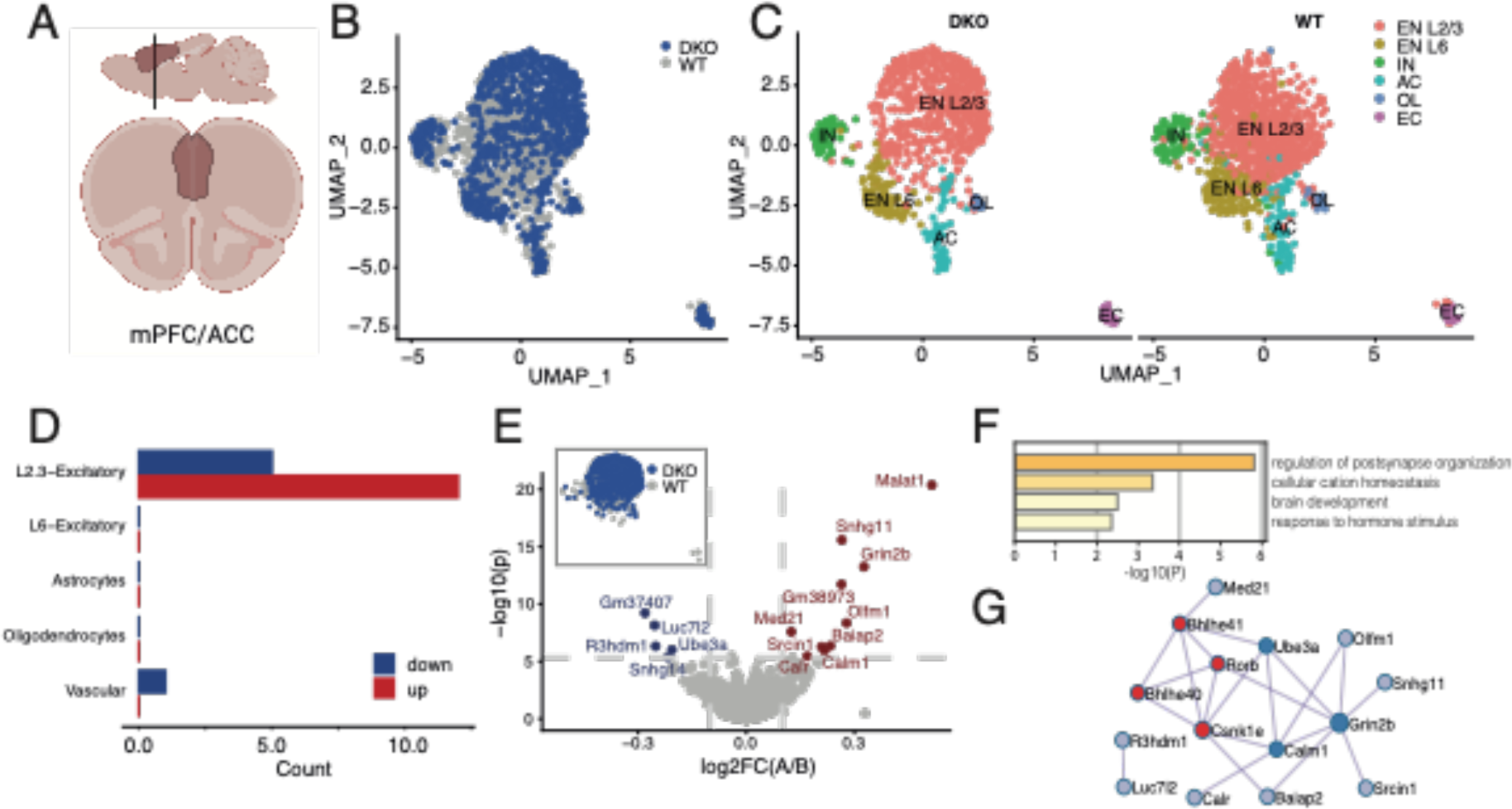
Layer 2/3 excitatory neurons in the mPFC of BHLHE40^-/-^/41^-/-^ mice show transcriptional changes in genes associated with the post-synapse. (A) The anterior cingulate cortex (ACC) part of the medial prefrontal cortex (mPFC) was selected as region of interest and isolated at Zeitgeber Time (ZT) 4, at circadian trough of BHLHE41 expression for single-nucleus RNA sequencing (snRNAseq) to identify constitutively deregulated genes. (B) UMAP dimension plot of the integrated datasets from wildtype (WT) and BHLHE40^-/-^/41^-/-^ (DKO) shows successful integration. (C) Six major cell populations, shown in distinct colors, were identified in both WT and DKO tissue samples. (D) Barplot of numbers of significantly deregulated genes between the genotypes in all cell clusters reveals a selective response in layer 2/3 cells only. (E) A Differential expression analysis between WT and DKO Layer 2/3 excitatory neurons (EN L2/3) identified a small set of differentially expressed genes. The gray lines indicates a threshold of 0.1 average log2-transformed fold-change (log2FC) and the Sidak-corrected significance threshold of the-log10-transformed p-value, assuming independent tests. (F) Gene ontology clusters enriched in the differentially expressed gene (DEG) set find with MetaScape^1^ hint to a postsynapse-related mechanism. (G) The top genes identified in the human GWGAS (red) form a protein-protein interaction (PPI) network with the DEGs identified in ACC-derived EN L2/3 (blue) with Grin2b, Calm1, and Ube3a (dark blue) at its center. EN L2/3: excitatory neurons layer 2/3 (upper layer); EN L6: excitatory neurons layer 6 (deeper layer); IN: inhibitory neurons; AC: astrocytes; OL: oligodendroglial cells; EC: endothelial cells

**Figure S11.**
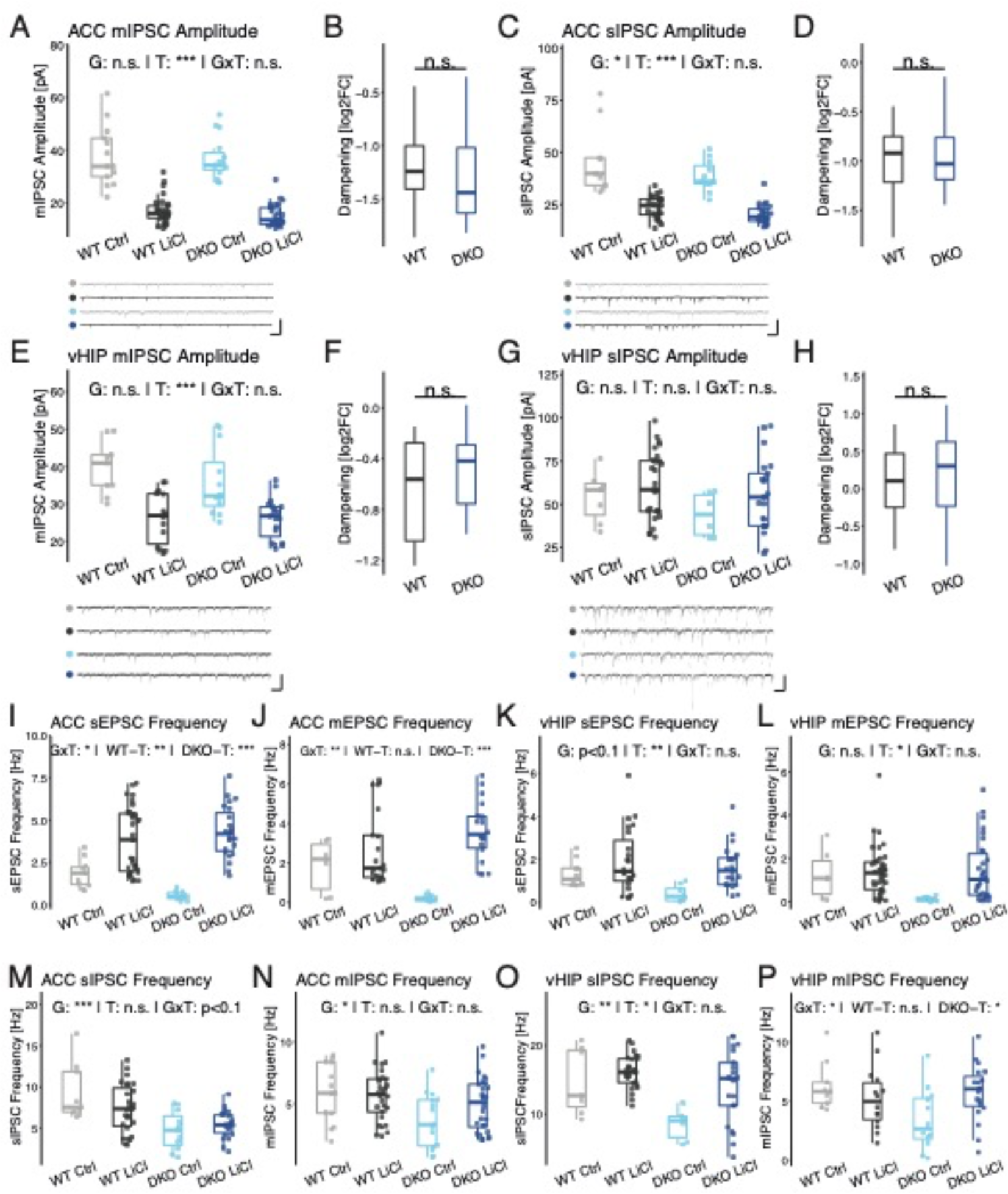
Extended data from whole-cell recordings in ACC and vHIP. Whole-cell voltage-clamp recordings in layer 2/3 pyramidal cells in the anterior cingulate cortex (ACC) area of the medial prefrontal cortex (mPFC) and pyramidal cells in the CA1 region of the ventral hippocampus (vHIP). (A-H) Average miniature and spontaneous inhibitory post-synaptic potential (mIPSC & sIPSC) amplitudes with example traces (scale bars x: 0.2 s & y: 0.1 nA) and the log2-transformed fold change of the respective amplitudes in LiCl compared to Ctrl mice. (I-L) Average miniature and spontaneous excitatory post-synaptic potential (mEPSC & sEPSC) frequency. (M-P) Average mIPSC and sIPSC frequency measured in ACC and vHIP. * P < 0.05; ** P < 0.01; *** P < 0.001; n.s. not significant; P-values refer to univariate two-way ANOVA with Type 2 sum-of-squares; simple effects were tested in a similar but unifactorial ANOVA procedure; WT: wildtype mice; DKO: Bhlhe40/41 double-knockout mice; Ctrl: vehicle control; LiCl: lithium-treated; G: genotype main effect; T: treatment main effect; GxT: interaction effect; simple T: simple effects; DKO-T: DKO simple treatment effect; WT-T: WT simple treatment effect.

**Figure S12.**
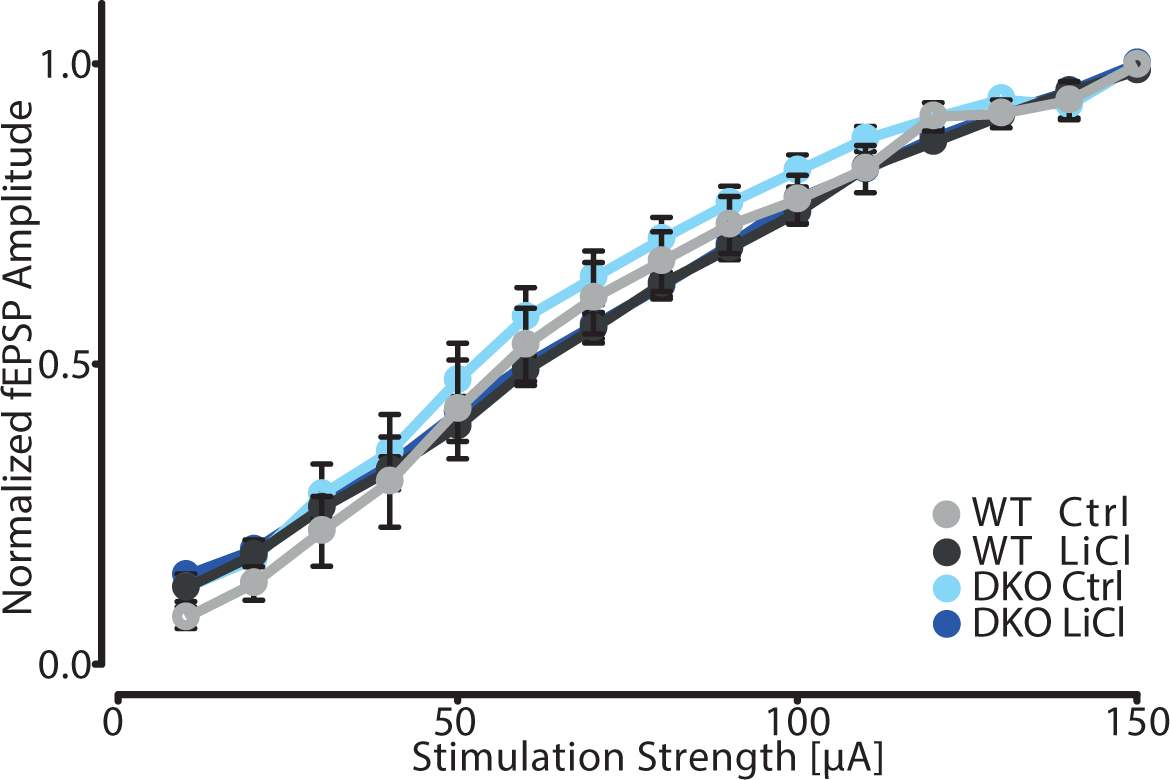
Extended Data on LTP Recordings in the CA1 Region of the Hippocampus. Input output (I/O) relationship of the mean (+/-SEM) normalized field excitatory post-synaptic potential (fEPSP) measured in the CA1 region plotted against the stimulation current injected at the Schaffer collaterals in the stratum radiatum at the CA3/CA1 junction did not show any differences in basic synaptic transmission between groups WT: wildtype mice; DKO: Bhlhe40/41 double-knockout mice; Ctrl: vehicle control; LiCl: lithium-treated

**Figure S13.**
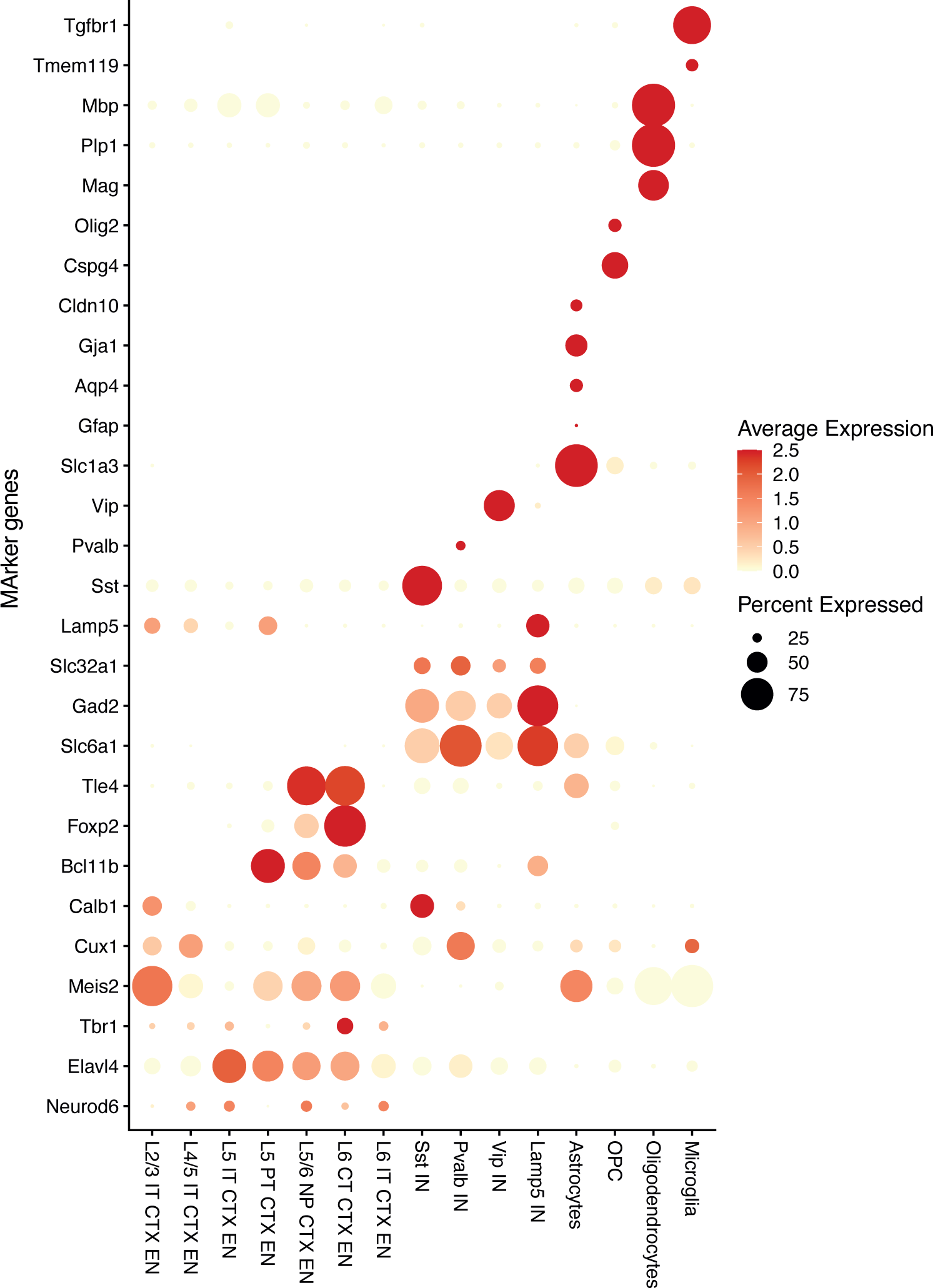
**Dotplots of selected cluster marker genes**. Normalized average expression is shown in color scale, abundance of expressing cells in percent as dot size.

**Figure S14.**
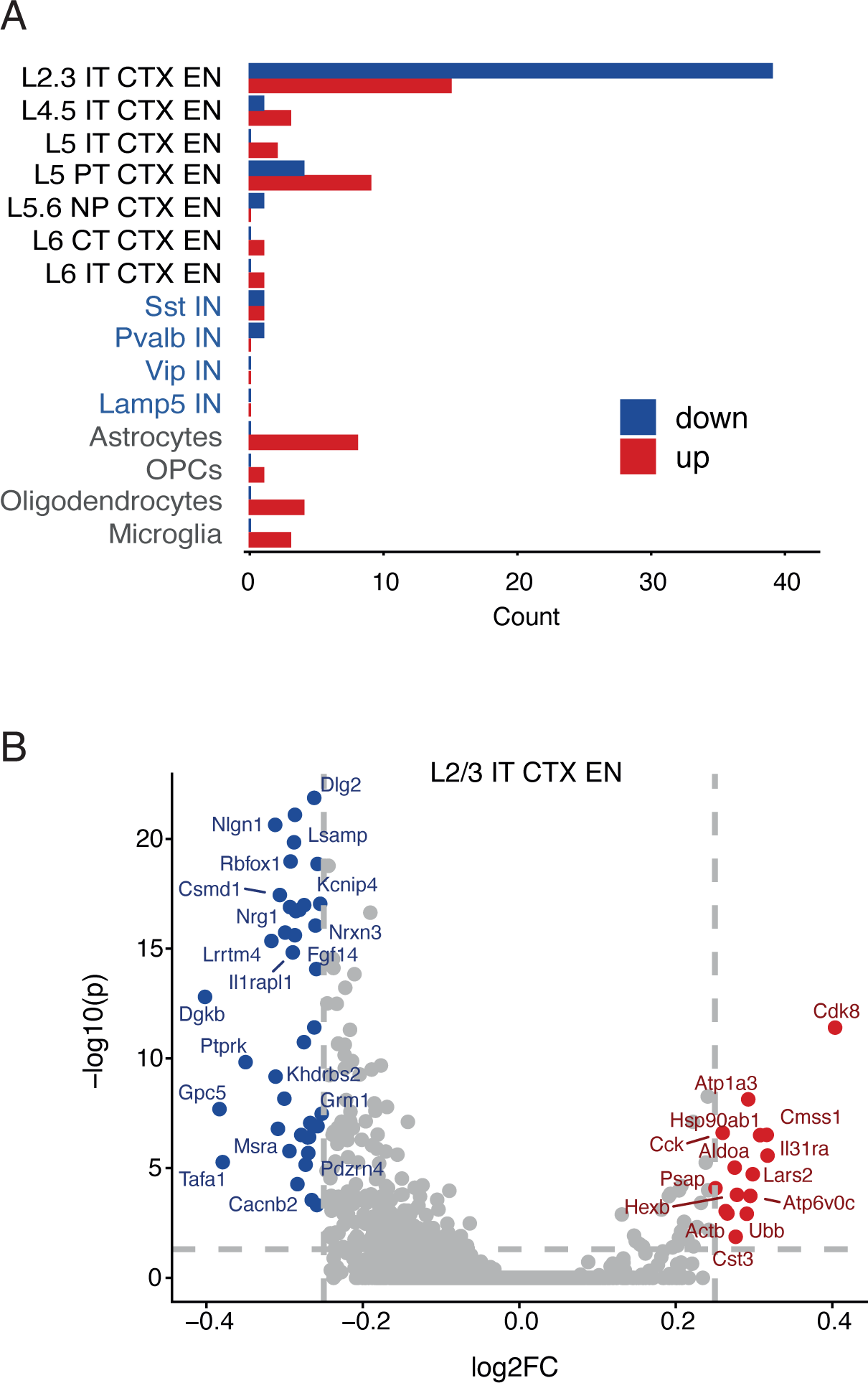
Differential gene expression analysis in wt and DKO treated with lithium by single cell RNAseq. (A) Barplot of numbers of significantly deregulated genes between the groups reveals a highly selective response in layer 2/3 excitatory neurons compared to all other cell types. (B) Volcano plot of deregulated genes in layer 2/3 excitatory neurons. On the Y axis the-log transformed adjusted P-value is shown and on the X-axis the average log2-transformed fold-change is plotted at the indicated thresholds (dashed lines).

**Figure S15.**
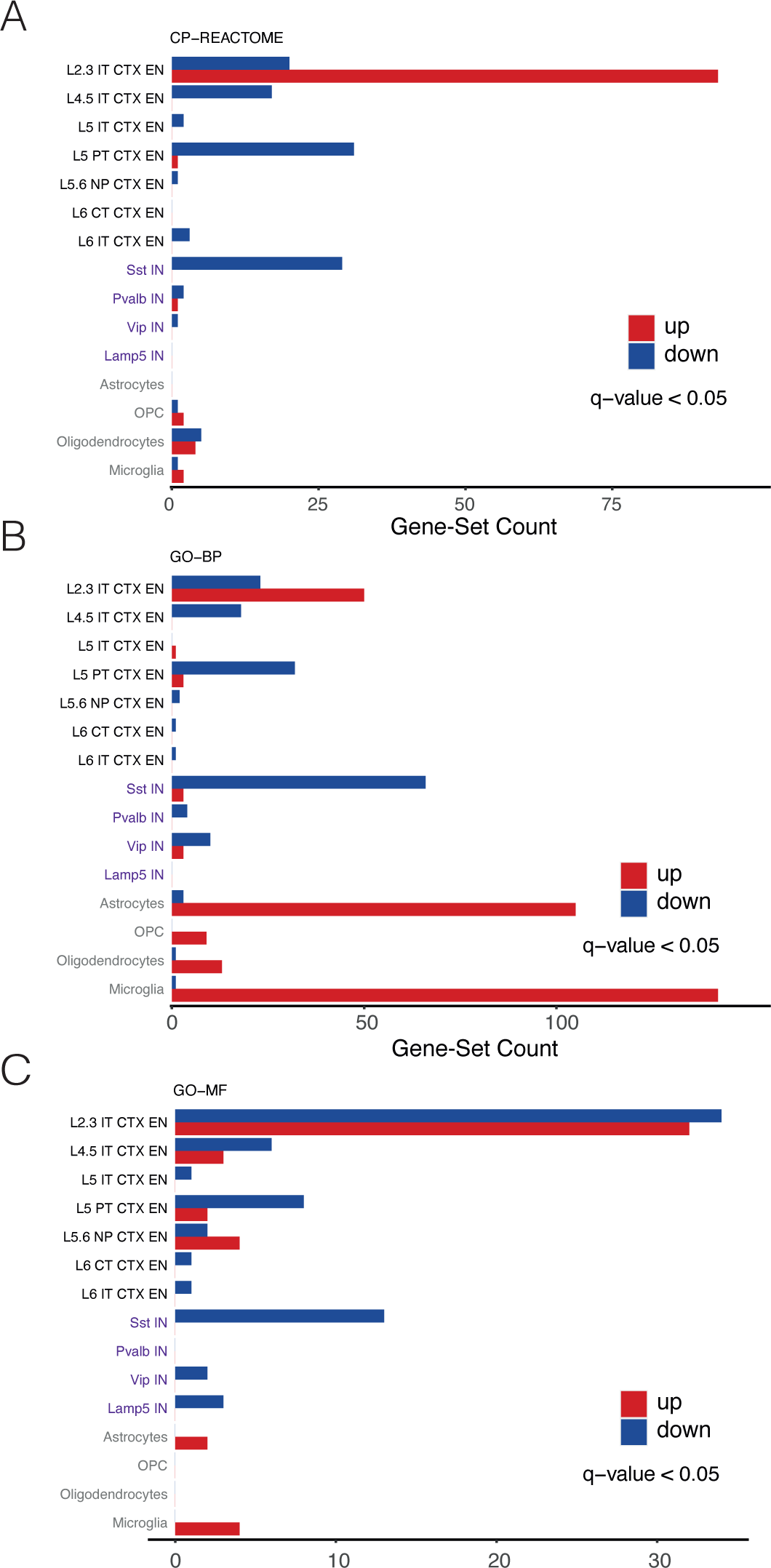
Differentially regulated gene-sets in wt and DKO treated with lithium identified with GSEA. (A-C) Barplot of numbers of significantly deregulated gene-sets (corrected q-val < 0.05) between the groups reveals a prominent response in layer 2/3 excitatory neurons in the pathway collection from reactome (A), gene ontology biological process (GO-BP) (B), and the gene ontology molecular function (GO-MF) (C) subcollections.

**Figure S16.**
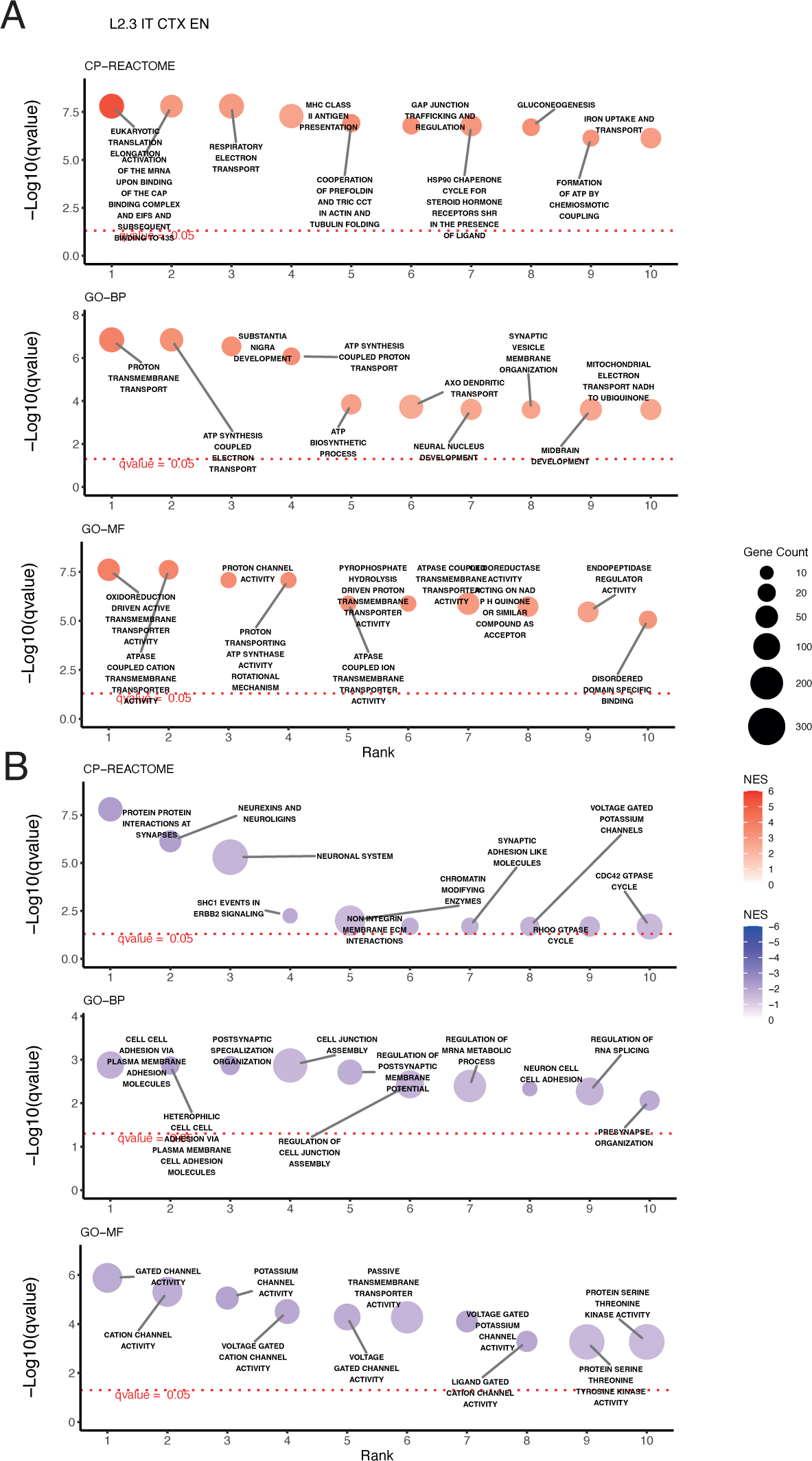
Top deregulated gene-sets in layer 2/3 excitatory neurons from wt and DKO treated with lithium identified with GSEA. (A) Ten most significantly upregulated and (B) downregulated gene-sets from the pathway databases reactome (top), the gene ontology biological process (GO-BP) (middle), and the gene ontology molecular function (GO-MF) (bottom) subcollections. Upregulated gene-sets are mainly associated with metabolism and mitochondrial ATP-synthesis. Down-regulated gene-sets are mainly associated with the synapse and cation-channel activities.

**Figure S17.**
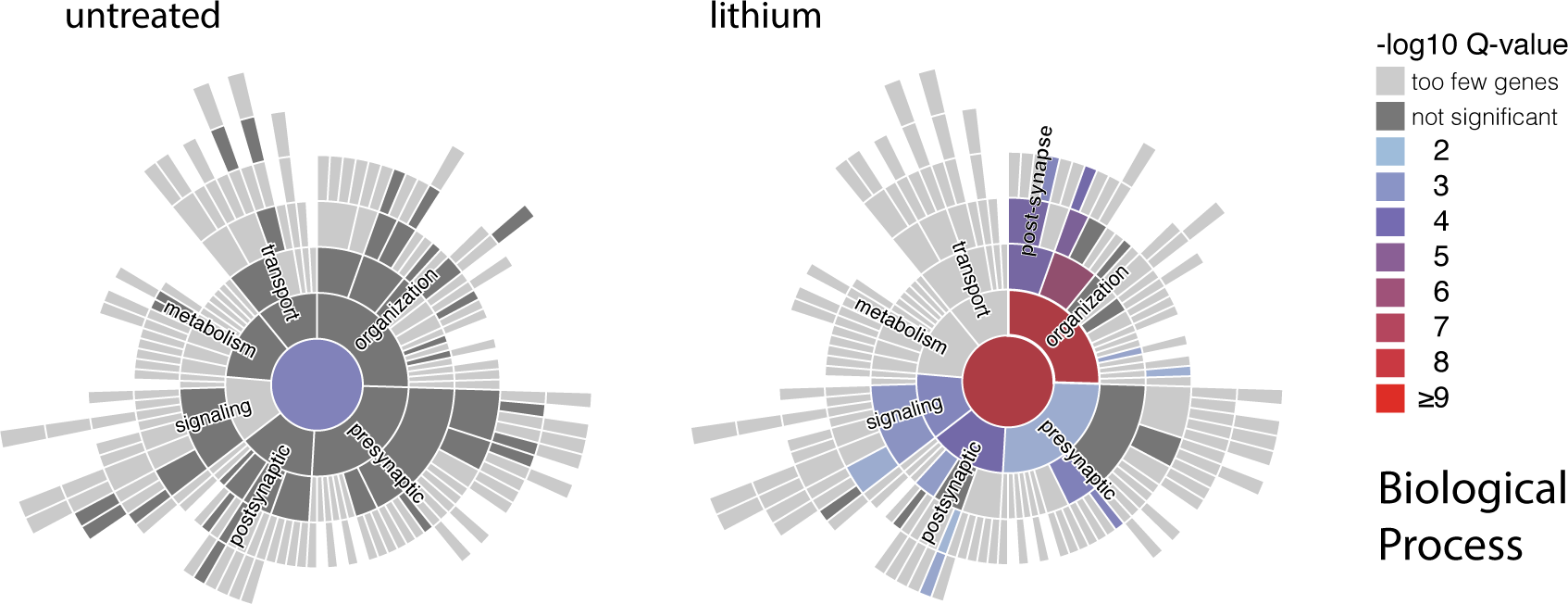
SynGO enriched gene-sets in layer 2/3 excitatory neurons from untreated and lithium treated wt and DKO scRNAseq ACC samples. Enrichment of synaptic gene-sets from the SynGO reveals most significant effects in lithium treated animals in biological processes associated with synaptic organization and post-synaptic functions.

## Online Methods

### Human Genetics

#### Definition of circadian gene sets

Core clock gene sets: genes centrally involved in the cellular time-keeping mechanism ^1–3^. Since it is difficult to define a universally accepted consensus list of core clock genes, we opted for three different gene sets ranging from a more restrictive (“Coreclock minimal”) to a more relaxed (“Coreclock extended”) definition.

Clock modulator gene set: *Clock modulators human* includes genes that serve as likely regulators of the core clock based on a genome-wide small interfering RNA (siRNA) screen in a human cellular clock model ^4^. The primary hit list included i) double hits (siRNA) for high amplitude, short period and long period, and ii) single hits for high amplitude and short period.

Circadian oscillating/output gene sets: genes that exhibit oscillatory expression but make no known contribution to the core time keeping mechanism. The *Circadian 6 mouse tissues* gene set was based on those genes showing a circadian oscillation expression pattern in at least 6 different mouse tissues ^5^. The *Circadian 4 CNS tissues (CircaDB)* gene set was based on the data available in CircaDB (http://circadb.hogeneschlab.org/) and included genes showing a circadian expression pattern in at least 4 mouse CNS tissues^6^. The *Circadian brain human* gene set included the top cyclic genes in the human brain across six different brain areas ^7^. Likewise, the *Circadian prefrontal cortex human* gene set represented the top 50 clock-regulated genes in the human prefrontal cortex (Brodmann areas 11 and 47) ^8^. A similar selection of the top 50 clock-regulated genes was done for the dorsolateral prefrontal cortex. *Circadian DL prefrontal cortex human* and the striatum (*Circadian nucleus accumbens/caudate/putamen human)* ^9,10^. Finally, a gene set including the top 50 circadian genes most commonly found in different human tissues was also defined (*Human ubiquitous circadian gene* ^11^. See Table S1 for a list of the specific genes contained in each gene set.

A curated homology list (Mouse Genome Informatics, The Jackson Laboratory) was used to perform mouse-to-human gene mapping.

#### Target phenotypes

GWAS summary statistics available for European-origin populations for schizophrenia (SCZ) ^12^ ^13^, post-traumatic stress disorder (PTSD) ^14^, major depressive disorder (MDD) ^15^, bipolar disorder (BD) ^16^, autism spectrum disorders (ASD) ^17^, attention-deficit/hyperactivity disorder (ADHD) ^18^, lithium response defined as continuous or dichotomous trait (ConLiGen) ^20^, and chronotype as positive control ^21^. For the independent replication study, lithium response GWAS summary statistics from a Swedish cohort was used ^22^. The sample size of each study is shown in Table S1.

#### Lithium response: samples and phenotypes

The international Consortium on Lithium Genetics (www.ConLiGen.org) has recruited 2,563 BD patients treated with lithium and includes 22 sites across four continents (Europe, America, Asia and Australia). Only European-origin ConLiGen samples (N = 2,343) were used for gene-set association (summary statistics) and polygenic analyses (N = 2,153). The “Retrospective Criteria of Long-Term Treatment Response in Research Subjects with Bipolar Disorder” (Alda scale) ^23^ was used for the evaluation of long-term treatment response to lithium. This scale was designed to assess long-term response, while considering confounding factors. The A score (range 0 – 10) quantifies symptom improvement over the lithium treatment period. The A score is then weighted against five criteria (B score) that assess confounding factors (disease course, compliance, and add-on treatments) each scored 0, 1, or 2. The total score is derived by subtracting the total B score from the A-score. Dichotomous response criteria were defined with a total score ≥ 7 for “responders” (< 7 “non-responders”). The continuous response was defined using A score but excluding subjects with a total B score > 4 ^24^.

In the replication Swedish cohort of BD patients, lithium response was defined on the basis of one subjective and one objective assessment ^22^. Subjective assessment of lithium response in participants who were treated for at least 1-year (N=1,120) was measured by means of a telephone interview, using a standardized interview protocol. Patients who reported complete remission on lithium were considered “responders”. Objective assessment of lithium response was based on recurrence data of patients (N=942) at yearly longitudinal follow-ups extracted from the Swedish Quality Register for BD (BipoläR). Responders were defined as having no mood episodes over at least 1 year of follow-up.

#### Gene set analyses

The European panel of 1000 Genomes Project, phase 3 was used as reference for all processing/analysis steps. NCBI build 37.3 genomic annotation was used for gene locations.

MAGMA ^25^ and INRICH ^26^ were used to carry out competitive gene set analyses. Both methods were chosen because of their good performance in keeping a low type-1 error rate and dealing with confounding factors like gene size, gene density and linkage disequilibrium (LD) between genes ^27^. Together they provide complementary approaches because each method captures different facets of the association signal ^27^. The built-in procedures for empirical multiple testing correction were not used. Instead, more conservative Bonferroni corrections were used for the set of traits and gene sets after enrichment analyses. Stratified LD Score Regression ^28,29^ was not used for enrichment analyses due to its low performance when using small annotations covering less than 0.5%-1% of the genome (https://www.medrxiv.org/content/10.1101/2021.03.13.21249938v1).

MAGMA (Multi-marker Analysis of Genomic Annotation, https://ctg.cncr.nl/software/magma) v1.09a was used for formal enrichment analyses using the summary statistics of the previously mentioned major psychiatric disorders, response to lithium treatment, and chronotype. After SNP-to-gene annotation, we performed gene-level analyses (± 10 kb) with the SNPwise-mean model. Gene-based P-values were obtained by combining the SNP P-values in a gene (± 10 kb) into a gene test statistic (mean χ2) corrected for linkage disequilibrium between SNPs. MAGMA competitive gene-set analysis was used within a regression framework to test whether the circadian-related gene sets were more strongly associated with the target phenotypes than other genes. For gene set competitive testing, linear regression analyses were conditioned to gene size, log(gene size), gene density, log(gene density), inverse mac, and log(inverse mac).

INRICH analyses (https://atgu.mgh.harvard.edu/inrich/) is a permutation-based competitive gene set analysis tool and takes a set of genomic regions (intervals) that are independently associated with a trait, and then tests for the enrichment of predefined gene sets. LD-independent associated genomic intervals for each trait were generated using SNP tagging (--tag-kb 1000--tag-r2 0.2) in PLINK (v1.9) ^30^. Intervals were calculated at three different P-value thresholds (P_TH_ = 0.00001, P_TH_ = 0.0001, P_TH_ = 0.001). The next step was to count the number of associated intervals overlapping with genes in the circadian gene sets. Interval overlap was limited to ±10 kb up/downstream of a gene. Random interval sets were generated 10,000 times (first-pass permutations), with each set approximately matching the associated intervals regarding the number of SNPs, overlapping genes, and SNP density per kb. The overlap of circadian gene sets was also determined in these random intervals to generate a distribution of the enrichment statistics for such target gene sets under the null hypothesis. Finally, the empirical P-value was calculated as the proportion of random replicates where the enrichment statistic is as large as that of the original interval set.

#### Data analysis and visualization

Additional statistical analyses and data management were carried out using R v3.5.0 and SPSS v25. Plots were generated using *ggplot2* or *lattice* R packages.

### Behavioral Animal Study

All animal experiments were carried out in accordance with institutional guidelines and regulations approved by the Regierungspräsidium Oberbayern, ROB Munich, Germany under the license ROB-55.2-1-54-2532-179-2016.

#### Animals and Husbandry

A wildtype colony of C57Bl/6NCrl mice was used for backcrossing. Chow and water were provided *ad libitum*, except for the IntelliCage system-based experiments for spatial learning. *Bhlhe40/41^−/−^* (DKO) and wildtype (WT) mice were no more than two to three generations apart from littermates and weaned after three weeks. Male and female *Arntl* (*Bmal1*) heterozygous and littermate wild-type control mice were used at an age of 8 weeks for behavioral analyses. Both, male and female mice were tested. The experiment mice were kept in type IV cages (Tecniplast 2000, 612 x 435 x 216 mm, 2065 cm²) in groups of litters of 12 to 16 mice. Experiments with young adult mice started at 8 weeks (Open Field Test) and ended at 19 weeks (Remote Fear Memory) of age.

#### Handling and General Experimental Procedures

Home cages were equipped with handling tunnels. All experiments outside of the IntelliCage system (TSE Systems) were conducted during the light phase, randomizing the trials across group to ensure a uniform distribution, and excluding circadian effects. For habituation, the home cage was placed in the test room for ten minutes before the actual experiment. Before and after each trial, the arena and other equipment in direct contact with test mice was cleaned with a dilute SDS solution, wiped dry, and disinfected with 70 % ethanol, unless stated otherwise. All behavioral test details have been described previously ^34^.

#### Lithium treatment

Starting three weeks before the first behavioral test, mice were treated chronically with 600 mg/L lithium chloride (LiCl) in the drinking water throughout the experiments. Water was available *ad libitum*. No significant differences in drinking behavior were observed between groups when monitored with the IntelliCage system.

#### Transponder Implantation

All experiment mice were equipped with an RFID chip for tracking in the IntelliCage and blinded identification throughout the experiments. Before the actual surgery, the neck region was shaved, cleaned with 70 % alcohol for disinfection, and the transponder (1.4 x 11 mm) was injected subcutaneously.

#### Behavioral Test

All behavioral tests, statistical analyses and visualizations were performed essentially as described in detail previously ^34^, and are described in detail below.

#### Camera Tracking

The following tests were done using the camera-based tracking software ANY-maze (Stoelting Europe): open field test, Y-maze test, tail suspension test, and fear conditioning test. In addition to the published standard fear conditioning protocol, the mice were placed in the original arena three weeks after the conditioning stage to assess remote context-related fear memory. All other tests were performed essentially as described in detail previously ^34–36^.

#### Prepulse Inhibition Test

For the assessment of prepulse inhibition of the startle response to auditory stimuli, we used the SDI Startle Response System and SRLab software (San Diego Instruments).

#### IntelliCage-Based Tests

For circadian activity tracking, spatial learning tests and the sucrose preference test, we used the IntelliCage system (TSE Systems). The device allows for home cage monitoring in social group and conditioning in four corners, each with a water bottle in a corner. The access to these water bottles can be denied in learning experiments by closing plastic doors, opening it only in case of “correct” nose pokes at the respective door. The tests were run as described ^34–36^ with the following minor modification: Activity was first assessed in a five-day period, during which the mice were first accustomed to the new environment (two days of free adaptation), the door opening (one day doors opened on visit), and to learn opening the doors by nose poking (two days nose-poke adaptation). After that, mice could access water in only one of the four corners for spatial positive reinforcement learning (two days place learning). Next the corner was changed pseudo-randomly for five days. Finally, one of the two bottles in each corner was filled with 4 % sucrose water for the sucrose preference test (one day).

#### Data Analysis and Visualization

Data were aggregated and processed using *FlowR* (XBehavior) as the user interface for the PsyCoP bundle. Additionally, R scripts were used to conduct all custom non-pipeline data analyses using *R 4.0.4* ^37^ in *RStudio IDE* (RStudio). Box and whisker plots and dimension plots were created with *ggplot2* ^38^ or *lattice* ^39^ R packages. Heatmaps were plotted with *pheatmap* ^40^. A multivariate linear model was fitted and used in a multivariate analysis of variance (ANOVA) from a Wilk’s lambda distribution using *car* ^41^. Univariate contrasts were derived from the same model. For univariate contrasts with a significant interaction, a unifactorial “simple-effects” ANOVA was computed to independently test the treatment effect for each genotype.

The dimensionality reduction procedure was described in detail earlier ^34^. In brief, missing values were imputed using the non-linear iterative partial least squares (NIPALS) algorithm in *ade4* ^42^. The resulting reconstituted matrix was used for canonical discriminant analysis (CDA) with *candisc* ^43^. CDA provides an optimal solution for group separation in phenotypic space, by finding linear combinations of single variables, called *canonical components,* similar to the *principal components* in a principal component analysis (PCA). Based on the CDA results of the collapsed factors, we generated a dimension plot with overlayed data ellipses, which include 75 % of the respective sample. A heatmap was created, showing the weights (*canonical coefficients*) of each dependent variable in the *canonical component* of each of the three terms (genotype, treatment, and interaction) of the multivariate linear model. Moreover, a CDA was computed for each of the two “simple-effects” models (DKO and WT).

Each variable was classified *a priori* in accordance with the Research Domain Criteria (RDoC) framework ^44^. The heatmap was sorted and grouped, accordingly.

### Single-Nucleus Transcriptomics

#### Sample Processing

For single-nucleus RNA sequencing (snRNAseq), mouse brains were isolated at zeitgeber time (ZT) 4, at the minimum of BHLHE41 expression ^45,46^, and kept in ice-cold phosphate-buffered solution (PBS; Gibco). Target tissue encompassing the anterior cingulate gyrus (bregma-0.1 – 1 mm) was isolated from 300 µm slices. Tissue from three mice was collected and pooled. The freshly isolated tissue was homogenized in nucleus isolation buffer (0.32 M sucrose, 3 mM Mg(Ac)_2_, 5 mM CaCl_2_, 10 mM Tris-HCl pH 8.1 with 0.1 % Triton-X100 freshly added; all Sigma) with a 2 ml Dounce-type tissue homogenizer. The sample was overlayed on top of a sucrose cushion (1.8 M sucrose, 3 mM Mg(Ac)_2,_ 10 mM Tris-HCl, pH 8.1, all Sigma) and centrifuged at 4 °C and 17,000 x g for 70 min. The supernatant was removed, the pellet washed with PBS and resuspended in resuspension buffer (PBS, 1 % bovine serum albumin (BSA) and 0.04 U/µL Protector RNAse inhibitor; all Sigma). The suspension was passed through 30 µm pre-separation filters (Miltenyi Biotec) and cells were kept in ice-cold resuspension buffer. Small aliquots of the samples were stained with trypan blue and Hoechst 33258 to determine nucleus density, integrity, and multiplet rate in an improved Neumann counting chamber. The nucleus density was adjusted to 2500 nuclei/L (±10 %). Nuclei aggregation rate was significantly below 10 % in all samples.

#### Library Preparation

Single-cell libraries were prepared according to manufacturer protocols (Illumina) and isolated using a ddSEQ Single-Cell Isolator (Bio-Rad) or the chromium controller (10x Genomics). Libraries were prepared following the manufacturer’s recommendations (SureCell 3’WTA (Illumina/BioRad) or Chromium Next GEM 5’ v2 (10xGenomics). Average fragment size, quality, and yield of the final libraries were assessed on a Bioanalyzer 2100 (Agilent). Libraries were sequenced at a loading concentration of 300 pM on a NovaSeq6000 instrument (Illumina) in paired-end mode.

#### Upstream Analysis of snRNAseq Data

The BCL files were demultiplexed with *bcl2fastq* (Illumina) and the sequence quality was checked with *FastQC* (v0.11.9). For 10x Genomics Next GEM libraries, barcode tagging, mapping and cell calling were done using CellRanger (v7.0; 10xGenomics). The reference genome was constructed from release 102 of the *Ensembl* annotation and the *GRCm38* mouse genome. For SureCell libraries, cell barcodes and unique molecular identifiers (UMIs) were isolated and tagged with *ddSeeker* ^47^. After tagging, the data were processed with *Drop-seq* tools (v2.3.0). A metabundle was created from the same genome assembly and annotation used for CellRanger. Corrupted cell barcodes were removed from the data. The reads were sorted by query name, and the data converted back to FASTQ for mapping. Read2 was mapped using *STAR aligner* (v2.7.5) ^48^ with default settings. Mapped data were merged with barcode-tagged data. The uniquely mapped reads were functionally annotated for counting. Digital gene expression data matrices were created for the 6000 most abundant cell barcodes in each sample, since no automated cell calling was implemented in the pipeline. The cut-off was chosen based on the knee plot made with *ddSeeker*’s *make_graphs.R*.

#### Downstream Analysis of snRNAseq Data

For downstream processing of snRNAseq data, we used the R package *Seurat v4* (v4.0.1) ^49^. After import as a dgc matrix, Seurat objects were created for each sample. Only cells with at least 200 features expressed and only features expressed in at least 3 cells were included in the object. Mitochondrial RNA content was determined by grepping for “^mt-” and hemoglobin contamination was quantified by grepping for “^Hb”. The cells were filtered for hemoglobin contamination of less than 1 % of the total UMI counts, a minimum of 500 (untreated) or 800 (lithium)different expressed features to remove both empty droplets and low-quality nuclei. Aggregates were identified and removed using the *scds* package’s combined approach ^50^.

#### Normalization and Integration of snRNAseq Data

The UMI count data matrices were normalized by the single cell transform procedure (SCTransform) ^51^. Both, mitochondrial and hemoglobin transcript contents were regressed out as indicators of RNA background. For integration of the lithium data, we used *Seurat’s* canonical correlation analysis (CCA) approach. For integration of the untreated dataset, we decided to shift to the reciprocal principal component analysis (rPCA) procedure implemented in *Seurat* with the *k.anchor* parameter set to 48, since we observed oversmoothing with CCA-based integration in that dataset.

#### Dimension Reduction and Clustering of snRNAseq Data

The first 30 principal components (PCs) were computed using the 3000 most variably expressed features. Using all 30 PCs, we performed similar nearest neighbor (SNN) graph-based clustering. We identified a total of 6 cell clusters in the untreated and 15 clusters in the lithium treated samples. Next, we computed a uniform manifold approximation and projection (UMAP) embedding ^52^.

#### Differential Expression Analysis

Differential gene expression was analyzed with *Seurat v4* ^49^. Features entering the analysis were filtered for a minimum of 1 % (untreated) or 10 % (lithium) expressing cells in the tested population. The filtered data were re-normalized to the lowest median UMI count sample but not integrated. The results were tested using the model-based analysis of single-cell transcriptomics (*MAST)* procedure ^53^ and P-values were Benjamini-Hochberg corrected. Differentially expressed features going into downstream analysis were filtered for a minimum average log2-transformed fold-change (avg_log2FC) of 0.1 (untreated) or 0.25 (lithium) as low coverage reduces the fold-change due to a high proportion of non-expressing cells.

#### General Enrichment Analysis and Protein Interaction Network Analysis

For gene annotation enrichment analysis and protein-protein interaction (PPI) network analysis, the Metascape platform was used ^54^. Differentially expressed genes (DEGs) from layer 2/3 neurons from the control samples were entered for custom analysis. Default parameters for annotation and membership were used. The background gene set parameter was set to the expressed features in the respective population and only selective gene ontology (GO) clusters were analyzed. For network analysis, the top 3 candidate genes, CSNK1E, BHLHE41, and RORB, as well as BHLHE41’s paralog BHLHE40, were added to the gene list and the network was computed using all databases combined.

Extended Protein-protein interaction maps with layer 2/3 excitatory neurons from the lithium samples were queried using the DGE genes as input using the Bisogenet plugin from Cytoscape which consolidates data from DIP, BIOGRID, HPRD, BIND, MINT and INTACT databases ^55,56^. Cytoscape network grafs were exported to Adobe Illustrator for final processing.

#### Gene Set Enrichment Analysis (GSEA)

The GSEA results were generated using an in-house developed wrapper for the clusterProfiler R package ^57^. The wrapper allows for selecting the desired gmt file/source and streamlining multiple analysis steps from data input to generation of results (tables and figures). The presented GSEA analysis was performed for GO terms (BP, CC, and MF) and Reactome pathways. The gmt data was obtained from MSigDb ^58^ using the msigdbr R package ^59^. The results were filtered for q-value ≤ 0.05 and normalized enrichment score (NES) > 0 for upregulated or NES < 0 for down regulated. ClusterProfilerWrapper Github repo: https://github.com/MNB-Lab/clusterProfilerWrapper

#### Enrichment Analysis of Synapse-Related Gene Ontology Terms

For a more specific enrichment analysis of synapse asspciated associated transcripts, the SynGO database for synapse related GO annotations was used ^60^. The input gene list was extended by relaxing the P-value adjustment and all expressed features in the respective cell population was used as background gene list.

### Reporter Assays

All reporter gene assays in primary cultured mouse cortical neurons were essentially performed as described previously ^61^. For data analysis in R, the mean signal and the corresponding standard error of each group was computed for plotting with ggplot2. For the boxplots and statistics, the area under the curve (AUC) was approximated using the rolling mean between neighboring datapoints for each sample’s stimulation peak and tested by two-way ANOVA using type 2 sum of squares and subsequent simple-effect (one-way) ANOVAs of the treatment effect of each genotype. Since a mean of means was tested and the small sample size of four, no test for normality was conducted.

#### Cell culture

Primary cortical neurons were isolated from brains of E15.5 C57BL6/NCrl WT and DKO mice. Cortical tissue was treated with a pre-activated papain suspension (Worthington) and gently dissociated by pipetting. 5 × 10^5^ Cells were plated in neurobasal medium supplemented with 5 % fetal calf serum (FCS), 2 % B27 serum supplement, and 1 % GlutaMax (all Gibco) on 35 mm plastic tissue culture dishes (BD Falcon) coated with poly-D-lysine (MW 70,000 – 150,000; Sigma).

On day *in vitro* (DIV) 1, plating medium was replaced with 1.5 mL culture medium (neurobasal medium, 2 % B27, 1 % GlutaMax). 500 µL culture medium was added on DIV5 and DIV11. The cultures were kept in a humidified incubator at 37 °C and 5 % CO_2_.

#### AAV production

AAV1/2 particles were produced following a previously published protocol ^61^. In brief, HEK293FT cells were transfected with the ESARE reporter AAV vector (pAAV-ESARE-Luc2A) and pFdelta6 helper plasmid. A mix of the capsid plasmids pH21 (serotype 1) and pRV1 (serotype 2) was used to produce AAV1/2 mixed serotype particles. Polyethylenimine (PEI) transfection reagent (Polyscience) was used for transfection. Cells were lysed and treated with benzonase (Sigma). The lysate was filtered through a 0.45 µm syringe filter (Millipore) and concentrated using an Amicon Ultra-15 centrifugal filter unit (100 kDa membrane cut-off; Millipore). Each AAV sample was digested with TURBO DNase (Invitrogen) to remove double-stranded copies of the viral genome and successfully packaged AAVs were released by proteinase K (Invitrogen) digestion and isolated by spin column-based purification (Macherey-Nagel). The titer was quantified by qRT-PCR with WPRE primers.

#### Live-cell luciferase assays

For ESARE assays, cultures were infected by adding virus stock to the culture medium on DIV1 (MOI 1000). On DIV5, lithium chloride in PBS or PBS alone was added with 500 µL medium (final concentration 1.5 mM). On DIV11, the supernatant was supplemented with D-luciferin for baseline recordings. After 24 hours, cultures were stimulated by adding 1(*S*),9(*R*)(−)-bicuculline methochloride (BIC) to a final concentration of 50 µM, silenced by adding a mixture of tetrodotoxin (TTX) and (2*R*)-amino-5-phosphonopentanoate (AP-V) at a final concentration of 1 µM (TTX) and 100 µM (AP-V), or mock treated by adding fresh medium (vehicle only). During the recordings all cell cultures were kept in a LumiCycle 32 apparatus inside a dry incubator (37 °C, 5 % CO_2_).

### Electrophysiological recordings

#### Whole-cell patch-clamp recordings

Slices were prepared as described previously ^62^. In brief, mice were decapitated, and the isolated brains transferred into ice-cold and pre-oxygenated artificial cerebrospinal fluid (aCSF; buffered solution of 1.25 mM Na_2_HPO_4_, 125 mM NaCl, 14 mM D-glucose, 2 mM MgSO_4_, 1.5 mM CaCl_2_, 2.5 mM KCl, and 26 mM NaHCO_3_, pH 7.4). 300 µm Slices were cut using a vibratome (VT 1200, Leica). Slices containing either the ACC or the vHIP were transferred into the incubation chambers with oxygenated aCSF (pH 7.4, aerated with 95 % O_2_, 5 % CO_2_, 32 – 34 °C) for 1 h recovery period, and then to room temperature aCSF, oxygenated with carbogen at a rate of 4 mL/min.

All whole-cell recordings were performed on pyramidal cells in the cortical layer 2/3 in ACC or pyramidal neurons in the CA1 region in vHIP, as described previously ^63,64^. Briefly, pyramidal neurons were identified by morphology. Access resistance was measured before and after recording using transient current responses to 5 mV hyperpolarization pulses. Patches with more than 10 % change in access resistance were excluded from downstream analysis. Additional exclusion criteria were leak currents of less than 150 pA, membrane resistance below 0.8 GΩ, or serial resistance of more than 20 MΩ.

Spontaneous excitatory postsynaptic currents (sEPSCs) were measured at a holding potential of −70 mV. sEPSC recordings in ACC were measured in the presence of 5 µM strychnine and 5 µM BIC. Recording pipettes were filled with recording solution (140 mM potassium gluconate, 0.5 mM Na_2_GTP, 4 mM Na_2_ATP, 10 mM EGTA, 1 mM CaCl_2_, and 2 mM MgCl_2_, buffered with 10 mM HEPES, pH 7.3).

Spontaneous inhibitory postsynaptic currents (sIPSCs) were measured in the presence of 10 µM 6-cyano-7-nitroquinoxaline-2,3-dione (CNQX, Sigma) and 50 µM AP-V. Here, the recording pipettes were filled with recording solution containing 0.5 mM Na_2_GTP, 4 mM Na_2_ATP, 10 mM EGTA, 140 mM KCl, 1 mM CaCl_2_, and 2 mM MgCl_2_, buffered with 10 mM HEPES (pH 7.2).

For mEPSC and mIPSC recordings, 0.5 µM TTX was added to the bath solution.

#### Long-term potentiation measurements

Long-term potentiation (LTP) recordings were performed as described earlier^65–67^. In brief, transverse hippocampus slices (300 µm) were cut in ice-cold and pre-oxygenated slicing buffer (10 mM glucose, 218 mM sucrose, 3 mM KCl, 0.2 mM CaCl_2_, 6 mM MgSO_4_, 1.25 mM NaH_2_PO_4_, and 26 mM NaHCO_3_). Slices were transferred to 32 °C warm aCSF oxygenated with carbogen and incubated for 1 h. For field potential recordings, Schaffer collaterals were stimulated in the stratum radiatum at the CA3/CA1 junction. Amplitude (baseline to peak) and slope (20 – 80 % level of the falling phase) of field excitatory post-synaptic potentials (fEPSPs) were quantified. The half-maximum response was defined as baseline fEPSP. Three trains of 1 s 100 Hz high-frequency stimulation (HFS) bouts in 20 s intervals were used for LTP induction. Every 20 s after the last train, the response to extracellular stimulation in the CA1 region was measured for 60 min. We quantified short-term potentiation (STP), defined as the mean response in the first 5 min after HFS, and LTP, defined as mean response in the last 10 min of measurement.

## Data Analysis

We quantified mean Amplitude and Frequency of mEPSC, sEPSC, mIPSC, as well as sIPSC in ACC and vHIP whole-cell recordings. Each variable was tested in a two-way ANOVA using type 2 sum of squares and, in case of a significant interaction, subsequent simple-effect (one-way) ANOVAs of the treatment effect for each genotype. Since a mean of means was tested, no test for normality was conducted. The same procedure was used for STP and LTP.

## Data Availability Statement

All data are available as Supplementary Tables, RNAseq raw data will be uploaded to GEO, codes are available via GitHub as indicated.

